# Multi-omics and functional analysis of a bioengineered vascularized pancreatic cancer model reveal an immunosuppressive and therapy-resistant niche

**DOI:** 10.64898/2026.03.05.709702

**Authors:** Giulio Giustarini, Ang Kok Siong, Pavanish Kumar, Germaine Teng, Basil Kuok Zi Xi, Chong Xiang Tan, Ragini Bhalla, Elisavet Kalaitsidou, Alicia Tay, Shanshan Wu Howland, Paola Cappello, Wei Wu, Jinmiao Chen, Salvatore Albani, Giulia Adriani

## Abstract

Pancreatic ductal adenocarcinoma (PDAC) is an aggressive disease characterized by therapy resistance and an immunosuppressive tumor microenvironment. To comprehensively characterize the complex stromal-immune cell interactions that drive PDAC aggressiveness, we applied an integrated multi-modal approach combining single-cell RNA sequencing, spatial transcriptomics, proteomics, immunofluorescence, and microfluidic-based functional assays to bioengineered spheroid models with increasing cellular complexity (up to four cell types) integrating human pancreatic cancer cells, pancreatic stellate cells, endothelial cells, and monocyte-derived macrophages. By incorporating vascularization within the OrganiX microfluidic platform, we enable studies of immune cell trafficking in vascularized tumors. Multi-omics phenotyping revealed coordinated molecular programs in our four-cell organotypic spheroid models, including enhanced hypoxic and glycolytic pathways, NF-κB activation, and ECM remodeling. Stromal and immune cells acquired tumor-associated phenotypes mirroring patient heterogeneity, including IL-1β+ macrophages, inflammatory cancer-associated fibroblasts (iCAFs), and antigen-presenting CAFs (apCAFs). The four-cell model exhibited superior clinical relevance, with gene expression signatures that correlated more closely with poor-prognosis patient cohorts and cancer hallmarks that were functionally validated through microfluidic-based assays demonstrating enhanced tumor invasion, angiogenesis, and therapeutic resistance. Finally, live imaging combined with transcriptomic readouts captures dynamic interactions between neutrophils and cancer cells in the vascularized microtumor, enabling direct observation of intravascular events relevant to metastatic dissemination. This integrated analysis demonstrated the recreation of a human-relevant aggressive PDAC niche, establishing a framework that bridges *in vitro* cellular crosstalk studies with patient-relevant therapeutic responses, offering a powerful translational tool for therapy development in PDAC.

## Introduction

Pancreatic ductal adenocarcinoma (PDAC) remains a clinical challenge as the third most lethal cancer, with low survival rates due to its late-stage diagnosis and limited treatment options [1]. Alarmingly, merely 24% of patients survive the first-year post-diagnosis [2]. The PDAC mortality arises not only from the absence of early symptoms but also from the poor efficacy of current therapeutic approaches [3]. The limited efficacy of the therapies is the result of testing on suboptimal preclinical models which fail to recapitulate the complex PDAC tumor immune microenvironment (TIME), leading to a lack of predictive power in therapy development [4]. The PDAC TIME is characterized by a highly desmoplastic and immunosuppressive milieu comprising diverse stromal and immune cell populations, extracellular matrix (ECM) components, and soluble factors that impact therapeutic efficacy [5].

Conventional two-dimensional (2D) cell cultures and murine models have failed to capture the heterocellular and ECM complexity that governs human PDAC progression, drug penetration and immune exclusion. Moreover, patient-derived models while clinically relevant, present challenges in scalability, reproducibility, and rapid implementation. Reliance on patient material complicates tissue acquisition, introduces considerable variability, and prevents standardization, thus limiting the development of robust and timely preclinical platforms. Together, these challenges leave an urgent need for refined three-dimensional (3D) human-relevant *in vitro* platforms that can bridge basic discovery with clinical translation [6].

To address these limitations, the past decade has seen considerable advances in the development of organoids and spheroids aimed at better replicating the TIME *in vitro* [7]. 3D culture systems have emerged as promising platforms to model solid tumors and predict therapy response [8–10]. In the context of PDAC, 3D *in vitro* models have shown the ability to emulate key features of the tumor complexity when stromal and immune cells were cultured together with cancer cells [11–13]. These studies have demonstrated that 3D *in vitro* PDAC models represent superior models for the investigation of molecular mechanisms due to their spatial cell organization and heterogeneity when compared to 2D cell cultures. Our recent work extended this complexity by creating a human heterotypic spheroid model including pancreatic cancer cells (PANC-1), pancreatic stellate cells (PSCs), endothelial cells (ECs) and peripheral-blood monocytes that developed a hypoxic core, a compressed endothelial network, and a cytokine signature characterized by IL-6, CXCL10, and CCL2, mirroring the immunosuppressive milieu found in patient tumors [13]. Notably, the integration of organoids and spheroids into organ-on-a-chip systems has improved control over microenvironmental factors, enhancing the physiological relevance and predictive power of 3D *in vitro* solid tumor models for studying drug delivery and immunotherapy as we have previously reviewed [10]. Despite these remarkable advances in 3D modelling, a gap in the current literature is the lack of integrated multi-omics and functional analyses capable of comprehensively dissecting cell–cell communication and molecular crosstalk in a fully *in vitro* heterocellular context.

Therefore, in the present study, we provide an integrated molecular and functional characterization of 3D *in vitro* PDAC spheroid models by combining single-cell RNA sequencing (scRNA-seq), proteomics, immunofluorescence (IF) followed by vascularization within the OrganiX microfluidic platform to enable 3D functional assays and spatial transcriptomics [14]. Through this integrated multi-modal approach to dissect the tumor immune microenvironment, we demonstrated that our four-cell organotypic spheroid model shapes a functional niche, reproducing the cellular interactions of the aggressive, immunosuppressive and therapy-resistant PDAC TIME more effectively than monoculture or simpler co-culture spheroid models. Specifically, we show that stromal and immune cells within the four-cell spheroids undergo phenotypic differentiation into tumor-associated macrophages (TAMs), and cancer-associated fibroblasts (CAFs), as defined in patients. The gene expression signature of the four-cell spheroids also correlates more closely with poor-prognosis of PDAC patient cohorts, demonstrating enhanced prognostic value compared to less complex models.

## Results

### Heterocellular spheroids establish a tumor–stroma–immune niche

We formed spheroids over 7 days by hanging drop technique as previously reported [13], starting with PANC-1 monoculture (C) spheroids and incrementally adding human PSCs to obtain bi-culture (CS) spheroids, human umbilical vein endothelial cells (ECs) for tri-culture (CSE) spheroids and peripheral blood mononuclear cells (PBMCs)-derived monocytes for four-cell (CSEM) organotypic spheroids (Figure 1A). ScRNA-seq revealed that the cellular composition significantly influenced transcriptional profiles across spheroid types (Figure 1B). Unsupervised clustering identified 9 cell clusters in the spheroids (Figure 1C) with relative abundance varying significantly, especially upon co-culture with PSCs (Figure 1D, Supplementary data file S1). Six epithelial cell clusters were manually annotated based on canonical markers such as *EPCAM, KRT15,* and *KRT19* (Supplementary figure S1), while remaining clusters were identified based on Panglao DB as PSCs, macrophages/monocytes (Mo/Mac) and ECs (Supplementary data file S1). Comparative transcriptomic analysis of epithelial clusters revealed one of the clusters highly expressing EMT/metastasis-associated genes (*CALD1, AREG, VIM, ALDH1A1, CADM2, ANXA1, TMSB4X, RAB27B, SRGN*) together with the hypoxia-induced EMT driver *TP63* [15], thus this cluster was annotated as EMT while the remaining epithelial clusters annotated as Epi1-5 (Figure 1E, Supplementary data file S1 [14]. Epi1 dominated heterocellular spheroids (>66%) while Epi4 dominated monoculture spheroids (∼59%), and EMT cells comprised ∼10% across conditions (Supplementary figure S2). We then embedded CSEM spheroids with HLA-matched vasculature in OrganiX™ microfluidic inserts, establishing a vascularized model (Figure 1F) that enables spatial analysis and leukocyte infiltration studies. Spatial transcriptomics on vascularized CSEM spheroids identified 5 cancer cell clusters (CC1-5), 2 fibroblast clusters (Fibro1 and Fibro2) and 2 EC clusters (EC, EndMT) with distinct localizations (Figure 1G). Interestingly, the EndMT cluster showed enriched expression for both fibroblast and EC genes, representing mesenchymal transitioning ECs.

**Figure 1.**
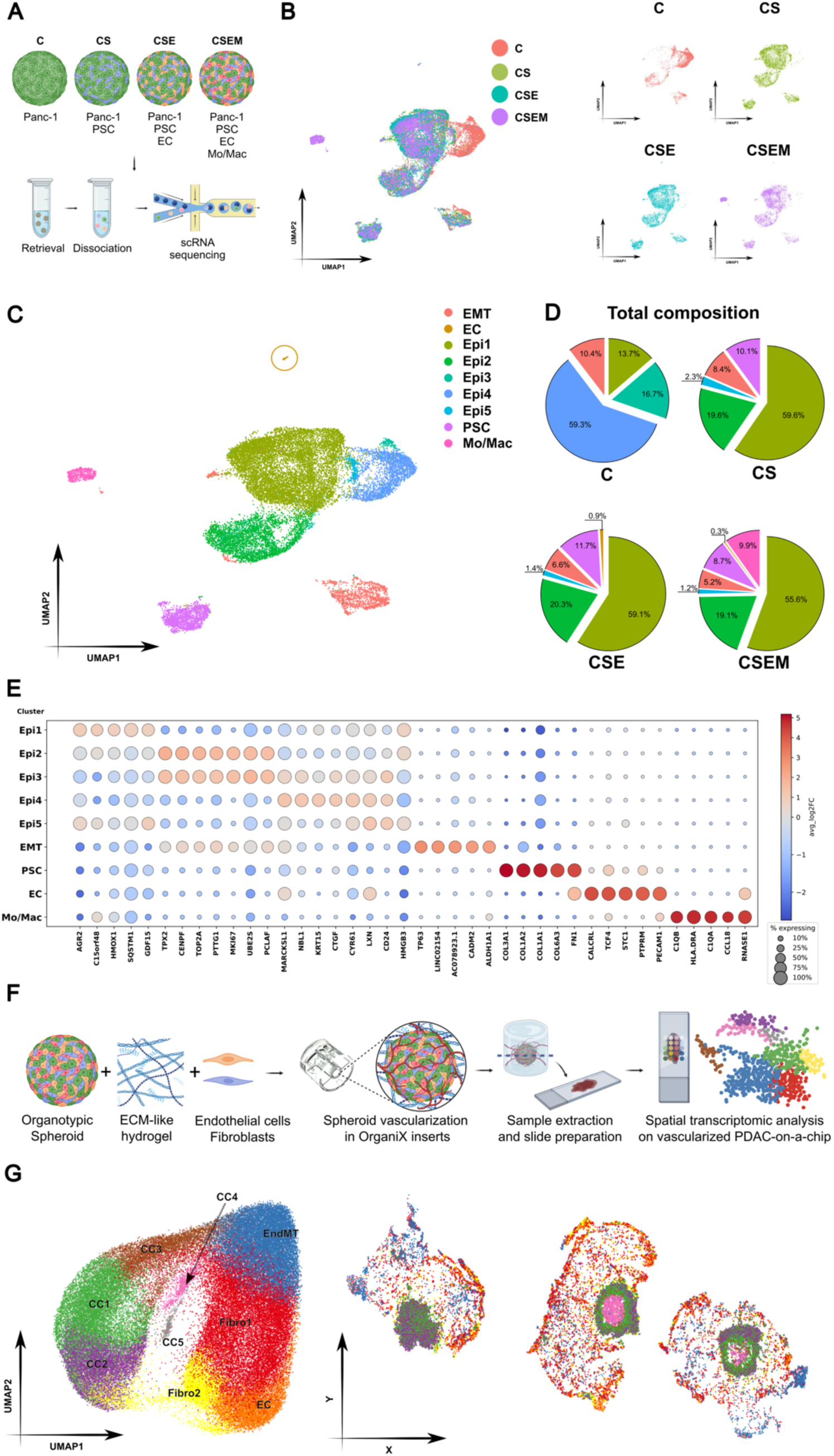
Single-cell and spatial transcriptomic characterization of pancreatic cancer spheroid and vascularized-on-chip models: microenvironment complexity drives shifts in cell composition. **(A)** Schematic of the experimental design showing four 3D pancreatic spheroid co-culture conditions: cancer cells alone (C), cancer cells with pancreatic stellate cells (CS), cancer cells with stellate and endothelial cells (CSE), and cancer cells with stellate, endothelial, and monocyte/macrophages (CSEM). Spheroids were retrieved, dissociated, and analyzed by scRNA-seq. Schematic created in BioRender.com. **(B)** UMAP visualization of the cell distribution for the spheroids colored by culture condition (left) and condition-specific UMAP plots (right). **(C)** UMAP visualization of integrated dataset colored by identified cell clusters, including six epithelial cell clusters (Epi1–Epi5, EMT), endothelial cells (ECs), pancreatic stellate cells (PSCs), and monocytes/macrophages (Mo/Mac). **(D)** Pie charts showing total cellular composition for each culture condition. **(E)** Dot plot of canonical marker expression across identified cell clusters. Dot size indicates the proportion of cells expressing the gene, and color represents the mean expression level. **(F)** Schematic of the workflow to generate vascularized pancreatic cancer-on-chip models, retrieval, and tissue sectioning for spatial transcriptomic analysis. Schematic created in BioRender. **(G)** Cell clustering (left) and spatial gene-expression map (right) after dimensionality reduction using scRNA-seq–derived differentially expressed genes.

### Spatial transcriptomics maps tumor and stromal programs in vascularized spheroids

To spatially map cellular programs within vascularized heterocellular spheroids, we performed stereo-seq on serial cryosections (S1–S3) of CSEM spheroids (Figure 2A). Cancer cell clusters distributed concentrically within the vascularized model: CC4 localized at the spheroid core, CC3, CC1 and CC2 positioned progressively toward the periphery, while CC5 comprised cells that had detached from the parent spheroid. GSEA revealed that CC5 upregulated interferon-α (IFN-α) signaling and the p53 pathway (Figure 2B), mirroring the EMT-like clusters observed in our spheroids before vascularization. Core-localized CC4 upregulated neuronal-associated genes, whereas both CC2 and CC1 were enriched for p53 signaling, with CC1 additionally scoring for TP53-regulated metabolic changes and hypoxia-activated genes consistent with oxygen diffusion limitation near the core (Figure 2B). Module score analysis using resected PDAC specimen gene signatures, showed that peripheral clusters exhibited basaloid transcriptional programs, while CC5 and peripheral cells associated with mesenchymal signatures; core cells showed significant similarities with the transcriptional profile of neural-like and neural-like progenitor cells (Figure 2C), that are transcriptional states associated with drug resistance and shorter prognosis in PDAC patients [16]. Pathway Enrichment Analysis (PEA) confirmed enrichment of lipid-homeostasis and glucose-metabolism pathways in CC1 (Figure 2D). Fibroblast clusters showed distinct spatial distributions with Fibro1 cluster spread in the gel of the vascularized model, and Fibro2 cluster concentrate mainly in proximity of the cancer spheroid (Figure 2A). Both fibroblast clusters showed ECM organization signatures, but Fibro1 was enriched for TGF-β signaling genes (*DCN, FBN1, FMOD, INHBA, FST, GREM1*) (Figure 2E), consistent with a myofibroblast-like state responsive to hypoxic signals. By contrast, Fibro2 upregulated prostaglandin synthesis genes (*PTGS1, PTGS2, AKR1C3, PTGES*) and TNF signaling via NF-kB genes (*CXCL1, CXCL3, IL6,TNFAIP3*) (Figure 2E), signatures of inflammatory CAFs (iCAFs) usually located adjacent to hypoxic areas [17].

**Figure 2.**
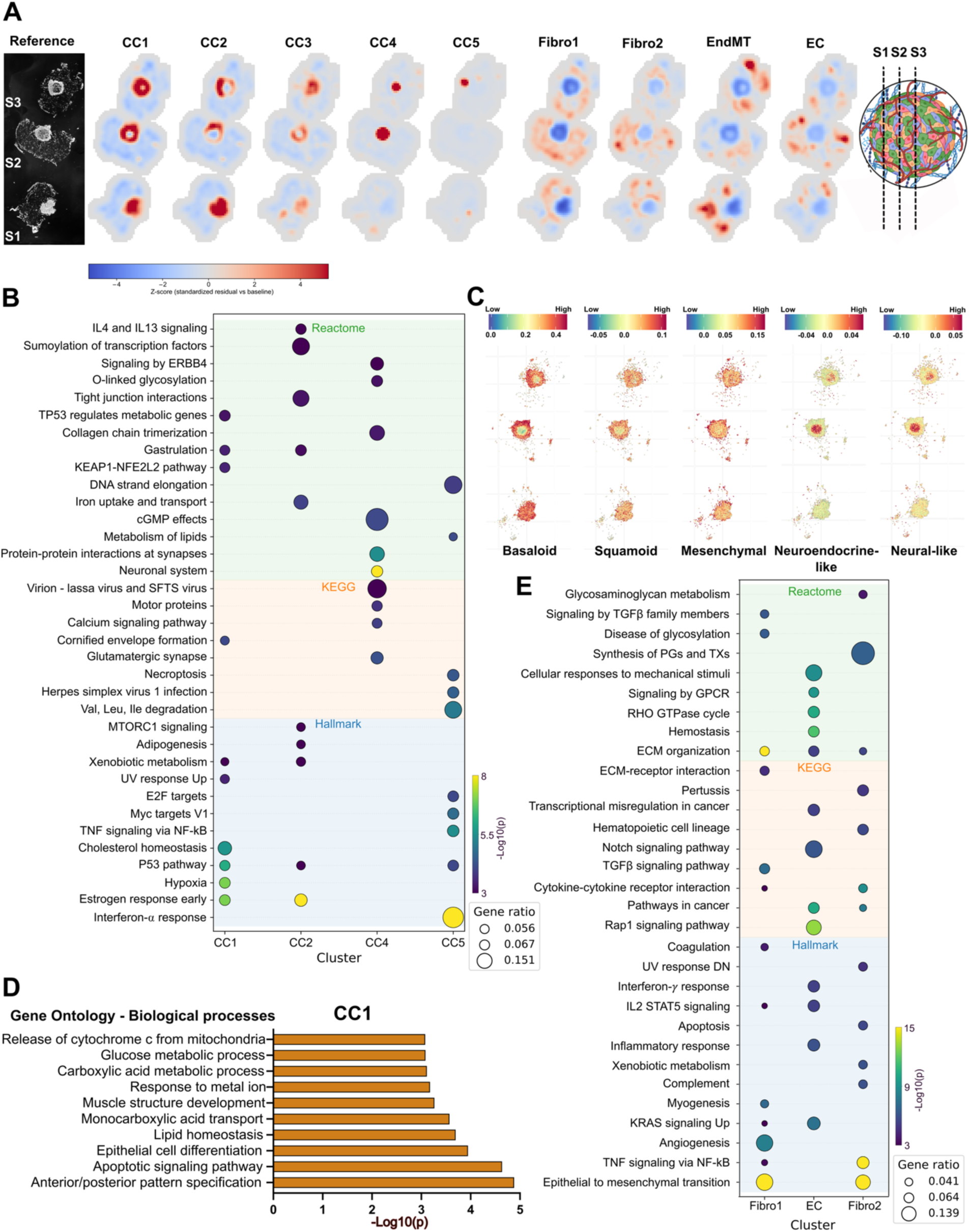
Spatial transcriptomics delineates tumor and stromal programs in vascularized heterocellular PDAC spheroids. (**A**) Reference images of three serial sections (S1–S3) and corresponding spatial maps of inferred cell-type enrichment for five cancer-cell clusters (CC1–CC5) and stromal/vascular populations (Fibro1, Fibro2, EndMT, EC). Enrichment is shown as a z-score (blue: low; red: high). The schematic (right) indicates the relative cutting planes through the spheroid. Schematic created in BioRender.com. (**B**) Bubble plot indicating gene set enrichment analysis of cluster-specific differentially expressed genes (Reactome, KEGG, Hallmark) across cancer-cell clusters. Dot size indicates gene ratio; dot color indicates significance (-Log_10_(p-value)). (**C**) Spatial distribution of epithelial transcriptional programs (basaloid, squamoid, mesenchymal, neuroendocrine-like, and neural-like) across sections; colors denote module score per cell. (**D**) Gene Ontology-Biological Process enrichment for CC1 highlighting metabolic and apoptosis-associated processes (bar length, -Log_10_(p-value)). (**E**) Bubble plot showing GSEA in stromal/vascular clusters (Fibro1, Fibro2, EC) illustrating dominant extracellular matrix, TGFβ-associated, angiogenic, and inflammatory/immune-response programs. Each dot corresponds to a pathway–cluster association; dot size represents the gene ratio and dot color encodes statistical significance (-Log_10_(p-value)).

### Heterocellular spheroids enrich hypoxic/iron-regulatory and proliferative cancer-cell states

To understand the biological implications of epithelial cluster variations across spheroid compositions, we performed PEA using the upregulated genes (-Log2FC > 0.5) from each epithelial cluster and identified the 5 most enriched Gene Ontology (GO) Biological Processes (BP) annotations for each epithelial cluster (Figure 3A). Epi1 cluster was associated with negative regulation of ferroptosis (*HMOX1, SQSTM1, FTH1*), intracellular iron ion homeostasis (*FTH1, FTL, HMOX1*) endoplasmic reticulum (ER) stress response (*HSPA5, AGR2, NUPR1*) and response to nutrient levels (*G6PD, HSPA5, UPP1, GDF15, AGR2*). In contrast, Epi2 and Epi3 clusters were enriched for chromosome organization and cell cycle regulation (*BIRC5, BUB1, BUB1B, CCNB1, CDK1, CDC20*), and centromere complex assembly (*CENPA, CENPE, CENPF, CENPI, NASP, DLGAP5*), indicating proliferative states. Interestingly, proliferative epithelial clusters showed distinct distributions across culture conditions, with Epi3 confined to monoculture spheroids and Epi2 emerging in co-culture spheroids (Supplementary Figure S2), with Epi2 cluster having 16.7% of cells in monoculture spheroids and Epi3 cluster ∼22-24% in heterocellular spheroids respectively, suggesting an increased rate of proliferation in these latter (Supplementary figure S2). The Epi5 cluster from heterocellular spheroids showed enrichment for IFN-α response (*IFITM1, IFITM3, IFITM2*), respiratory burst (*CYBA, CD55, CD24*) and actin filament organization (*ADD3*, *TMSB4X, PLEKHG2, CTNNA2, PFN1*), while Epi4 (predominant in C spheroids) associated with connective tissue development, and epithelial cell differentiation (*KRT8/9/10/19*) (Figure 3B). The EMT cluster exhibited associations with cytoplasmic translation, hormone response and “positive regulation of signal transduction by p53 class mediator” (gene ratio of 0.2; Supplementary data file S2) which could indicate a significant role of p53 in modulating EMT signaling pathways. The immunofluorescent staining confirmed a high expression of p53 (wild-type and mutant) in the cells of the CSEM spheroids which were located at the periphery of the spheroid or migrated from the primary tumor mass (Figure 3C). Spatially resolved transcriptomics identified an outer region of the vascularized CSEM spheroid enriched of cells with clear p53 signaling, G2M checkpoint and EMT transcriptional programs, while an inner lining of cells was enriched for ferroptosis inhibition genes (Figure 3D,E). The proteomic analysis comparing CSEM and C spheroids, corroborated the transcriptomic findings revealing that CSEM cancer cells were enriched in proteins related to DNA replication/repair (CDK1, MCM4, MCM7, RFC2, and POLE3), antigen processing and presentation (TAP1, TAP2, and TAPBP), and response to nutrient levels and starvation (ASNS, ATG7, LARS1) (Figure 4A,B), together with autophagy-related proteins SQSTM1 and TGM2 (Figure 4A,B). This profile indicates CSEM cancer cells adopting a highly stress-adaptive and immunologically active state that may support tumor growth and survival in a nutrient-limited, immune-competent microenvironment. Of note, C15orf48, GLA and DDT proteins, found to be upregulated in patient pancreatic cancer cells and exploited for vaccines [18,19], were significantly elevated in CSEM cancer cells versus C spheroids (Figure 4A,B).

**Figure 3.**
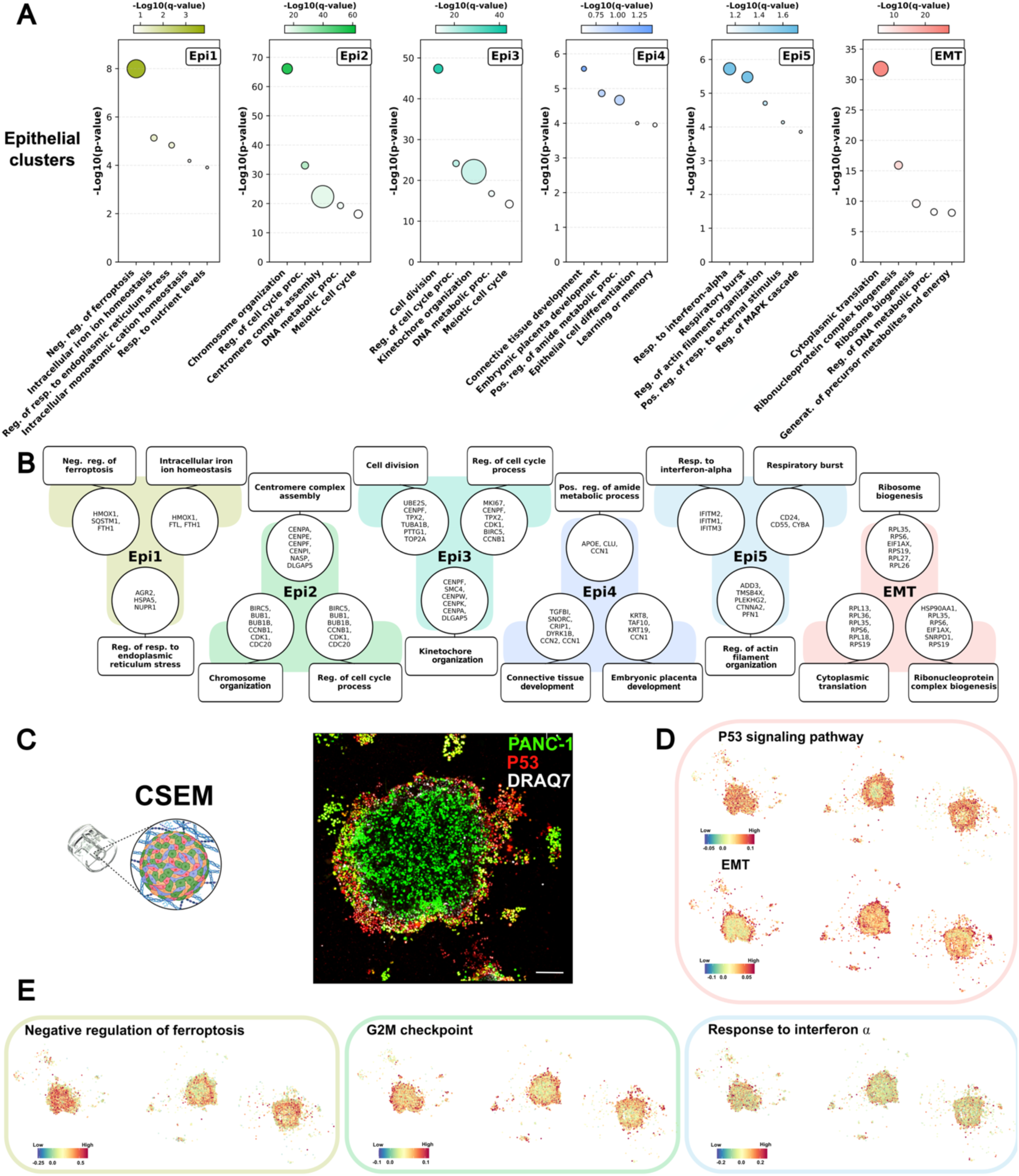
Functional and spatial heterogeneity in pancreatic cancer models: negative regulation of ferroptosis and proliferation define dominant epithelial clusters. (**A**) Gene ontology enrichment analysis of epithelial clusters (Epi1–Epi5, EMT) using upregulated genes (Log2(FC) > 0.5, adj. p-value > 0.05). Top: Bubble plots showing significant enriched pathways and biological processes, with color intensity representing q-values and bubble size representing the gene ratio (%). Bottom: Schematic representation of the top expressed genes and associated pathways for each cluster. (**B**) Spatial transcriptomic mapping of representative biological pathways across vascularized spheroid sections, illustrating enrichment of negative ferroptosis regulation, G2/M checkpoint activity, IFN-α response, p53 signaling, and EMT programs. (**C**) Schematic of spheroid cultured for 7 days in 3D within the hydrogel region of the OrganiX^TM^ inserts (left), and representative confocal images of C and CSEM spheroid models showing immunofluorescence validation of p53 expression (green: PANC-1; red: p53; blue: DRAQ7 nuclear stain). Scale bar: 200µm. Schematics created in BioRender.com. (**D**) Spatial visualization of p53 transcriptional signature in the CSEM spheroids. Color scale indicates module score for the mentioned gene set. (**E**) Spatial distribution of gene expression for the most significantly enriched biological process of each epithelial cell cluster in vascularized quadri-culture spheroids. Color scale indicates module score for the mentioned gene set.

**Figure 4.**
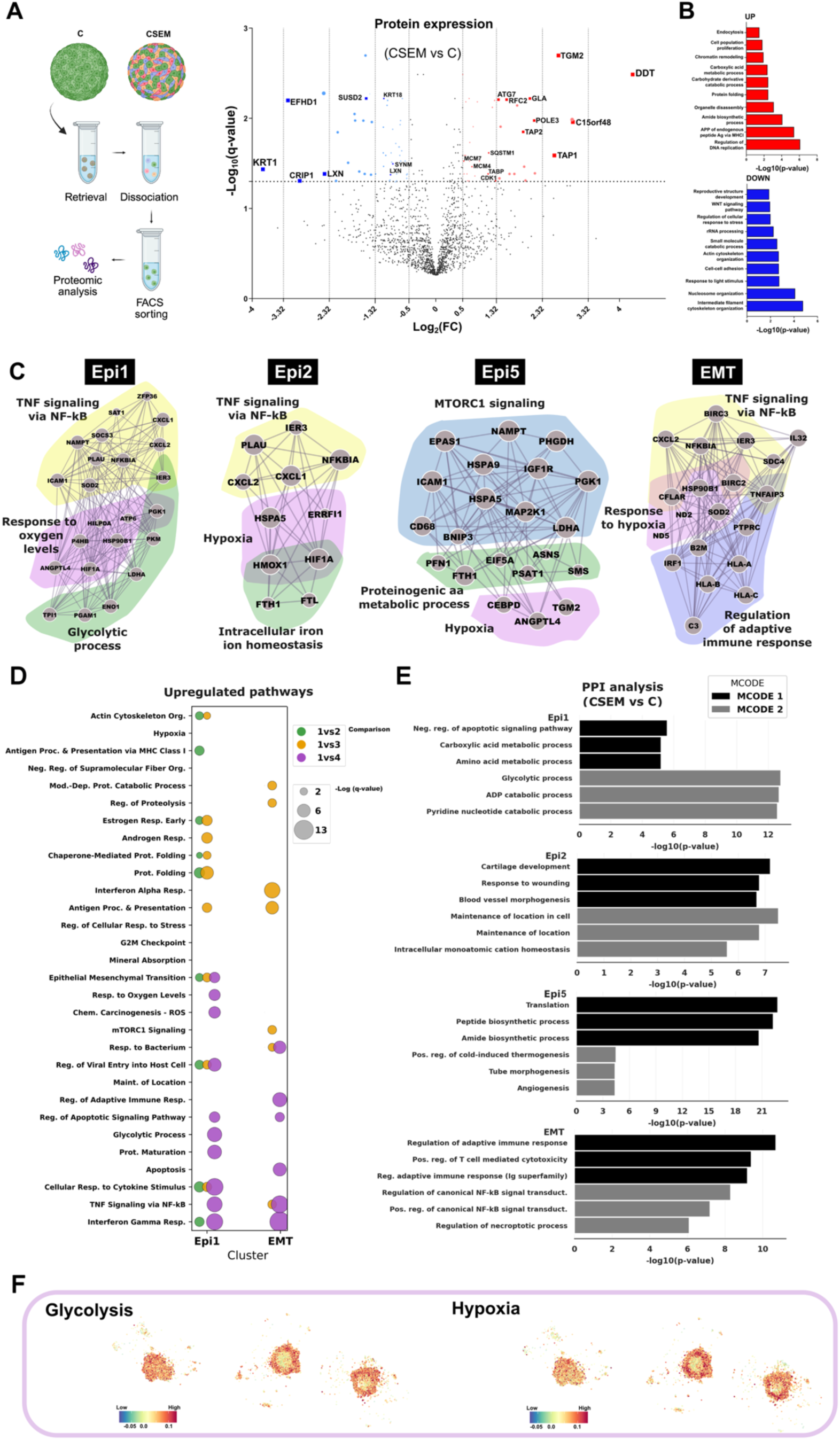
Comparative analysis of gene expression and protein-protein interaction networks in Epi1, Epi2, Epi5 and EMT clusters of monoculture spheroids versus heterocellular ones. (**A**) Workflow for the proteomic analysis using C and CSEM spheroids (left). Volcano plot of LC-MS/MS data from FACS-sorted GFP-PANC1 cells showing protein expression changes in CSEM versus C spheroids. Significantly regulated protein (FDR < 0.05) with variation greater than 1.5 folds (Log_2_(FC) > 0.585) are in red, and less than 1.5 folds (Log_2_(FC) < -0.585) in blue. N=2 biological replicates with technical duplicates. Schematics created using BioRender.com. (**B**) GO-BP pathway enrichment for strongly upregulated (Log₂(FC) > 1) and downregulated (Log₂(FC) < −1) proteins, with the top 10 terms for each direction shown as red and blue bar plot, respectively. (**C**) Protein-protein interaction (PPI) networks of upregulated genes associated with main annotations identified in each cluster. (**D**) Bubble plots of pathway and gene set enrichment analysis performed using upregulated genes (Log_2_(FC) > 0.5, adj. p-value > 0.05) of Epi1, Epi2, Epi5 and EMT cell clusters in heterocellular versus C spheroids. Color represents cluster, bubble size encodes -Log_10_(q-value). (**E**) Bar plots showing the first 2 MCODE algorithm results in Metascape using all DEGs (-0.5 Log_2_(FC) < DEG > 0.5 Log_2_(FC), adj. p-value > 0.05) for each epithelial cluster (as shown in figure). The three GO-BP annotations identified per MCODE result were used to generate the bar plots with their respective p-values. (**F**) Spatial distribution of glycolysis and hypoxia transcriptional programs in cancer cells across sections with color denoting program module score per cell.

### Negative regulation of apoptosis, NF-kB activation, hypoxia, and glycolysis characterize cancer cells in heterocellular spheroids

The transcriptomic analysis of epithelial cells identified in CSEM spheroids upregulated genes linked to apoptosis regulation, hypoxia, glycolysis, and TNF-NF-kB signaling (Figure 4C,D). Consistent with reports in many cancers including PDAC, NF-kB activation is known to support stemness, EMT, angiogenesis, and therapy resistance [20]. Epi1/2 and EMT clusters of the CSEM spheroids were enriched for hypoxia-associated annotations (Figure 4C,D), with additional signatures involving ferritin genes (*FTH1* and *FTL*) and IFN-induced transmembrane genes (*IFITM1-3*) (Supplementary data file S3), which are regulated upstream and downstream of the hypoxic pathway [21] and correlate with poor prognosis, EMT, angiogenesis and metastasis in preclinical *in vivo* and *in vitro* PDAC models [22,23]. Exclusively in the CSEM spheroids, Epi1 upregulated glycolytic pathways, while Epi5 cluster revealed enrichment for pyridine-containing metabolism, proteinogenic amino acid metabolism and mTORC1 signaling (Figure 4C). Notably, *PGK1* was among the most upregulated genes in both Epi1 and Epi5 clusters of the CSEM spheroids, consistent with macrophage-derived IL6-driven aerobic glycolysis [24], suggesting macrophage–cancer cell–induced metabolic flexibility (Figure 4C,D). Protein-protein interaction (PPI) analysis of DEGs between C and CSEM epithelia identified negative regulation of apoptosis and metabolic pathways (glycolysis, purine and pyridine catabolism) in Epi1, supporting a link between metabolic rewiring and apoptosis resistance (Figure 4E). Interestingly, the Epi2 and Epi5 DEGs highlighted their involvement in blood vessel morphogenesis/angiogenesis, suggesting an increased capacity of CSEM spheroids to sustain tumor vasculature formation (Figure 4E). Spatial transcriptomics of vascularized CSEM spheroid revealed glycolytic program intensifying in core-proximal regions, coincident with hypoxia signatures, suggesting a glycolytic, hypoxic area while aerobic glycolysis extending into more oxygenated peripherical regions (Figure 4F). Together, these data indicate that CSEM spheroids mimic critical malignant programs, which drive pathophysiological changes, PDAC aggressiveness and therapy resistance.

### Heterocellular spheroids recapitulate EMT-associated inflammatory and invasive pathways

The PPI analysis of DEGs in the CSEM EMT cluster identified two highly interconnected gene groups associated with positive regulation of canonical NF-kB activation and regulation of the necroptosis (Figure 4C-E). EMT cells showed higher expression of genes with anti-apoptotic and IFN/NF-κB regulating functions (*BIRC3, CFLAR, IRF1, ISG20, IFITM3, IER3*) together with mitochondrial ROS buffering superoxide dismutase 2 gene (*SOD2*), consistent with anoikis resistance and evasion of death-receptor–mediated apoptosis/necroptosis rather than ongoing cell death (Figure 4C, Supplementary data file S3). Additional upregulated genes suggest that in CSEM spheroids EMT cells potentially interact with immune cells and acquire higher migratory potential, as shown by the increased expression of the pan-leukocyte marker *PTPRC* (CD45) and invasive markers *IL32*, *SDC4*, and *LCN2* [25–28] respectively (Supplementary data file S3). The EMT cluster also upregulated semaphorin genes (Supplementary data file S1), proteins mediating invasion and migration of cancer cells in PDAC [29,30], and HLA class I molecules (*HLA-A, HLA-B, HLA-C*) (Figure 4C). These findings suggest that EMT cells of CSEM spheroids possess higher invasive potential, can initiate inflammatory cascades in the TIME, and based on their CD45 expression, may represent hybrid myeloid-cancer phenotypes as previously observed in other *in vitro* models [28]. Proteomic analysis validated these transcriptional changes, showing that cancer cells in CSEM spheroids downregulated proteins associated with intermediate filament cytoskeleton organization and cell-cell adhesion relative to monoculture (Figure 4A,B). Specifically, key cytoskeletal proteins KRT18, KRT1, and SYNM, were downregulated in CSEM spheroids (Figure 4A), indicating loss of polarity and adhesion, and gain of migratory and invasive traits [31]. Additional downregulated proteins included CRIP1, LXN, EFHD1, and SUSD2 (Figure 4A,B), which have been reported as decreased in pancreatic cancer or possessing tumor-suppressive, anti-invasive roles in PDAC and other malignancies [32–35].

### Heterocellular spheroids reprogram PSCs into WNT-activated, CAF-like states that remodel ECM and support vascularization

Genomic profiling of PSCs within heterocellular spheroids showed enrichment of canonical mesenchymal cancer programs, including ECM organization, blood vessel development and cell proliferation (Supplementary Figure S3). PSCs upregulated multiple interconnected ECM genes encoding collagens, metalloproteinases and enzymes mediating ECM cross-linking and degradation (Figure 5A), while ECs increased transcription of angiogenesis- and TGF-β–related genes, [36], and *COL4A1* (Supplementary data file S1), a marker of tumor-associated endothelial cells in PDAC [37], indicating coordinated stromal support for vascularization.

**Figure 5.**
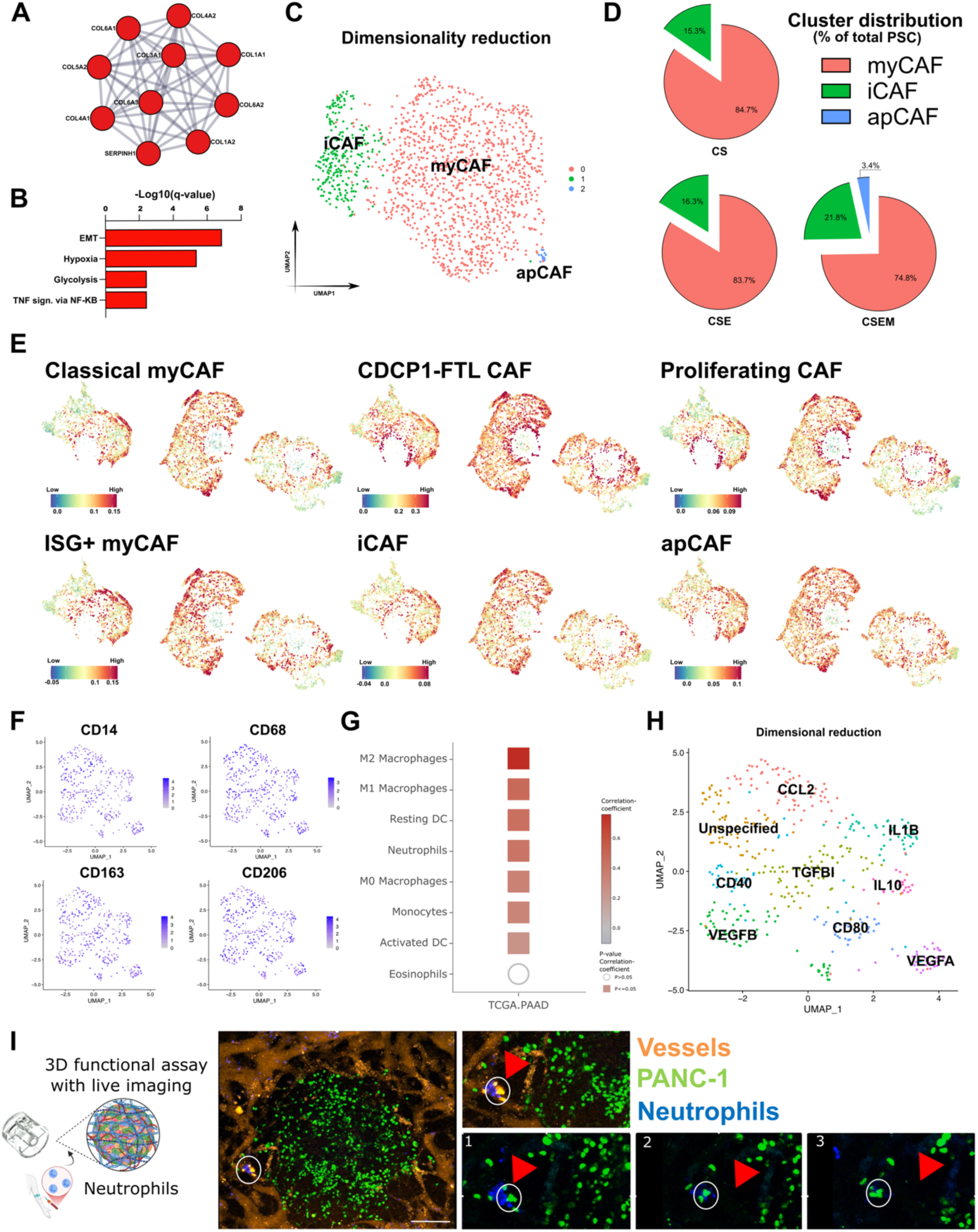
Stromal and myeloid remodeling in CSEM spheroids reveals CAF diversification, IL1B-high macrophages, and neutrophil–tumor intravascular interactions. (**A**) Network of upregulated genes associated with ECM organization and identified using the MCODE algorithm. (**B**) Bar graph of the Hallmarks annotations identified with the gene set enrichment analysis (GSEA) using upregulated genes in CSEM PSCs compared to CS PSCs. (**C**) UMAP visualization of PSC subtypes based on canonical CAF markers: myofibroblast CAFs (myCAF), inflammatory CAFs (iCAF) and antigen presenting CAFs (apCAF). (**D**) Pie charts showing the proportion of CAF subtypes within PSC population across spheroids with increasing levels of heterogeneity. (**E**) Spatial transcriptomic mapping of CAF associated with subtypes identified in PDAC patients, including classical myCAF, CDCP1–FTL CAF, ISG+ myCAF, proliferating CAF, iCAF, and apCAF. Colors denote program module score per cell. (**F**) UMAP visualization of the expression of canonical pro-inflammatory and anti-inflammatory monocyte/macrophage markers. Color scale represents the expression level of the gene. (**G**) Correlation of Mo/Mac gene signatures with immune cell types from the cancer genome atlas (TCGA) pancreatic adenocarcinoma (PAAD) dataset, indicating strong association with M2 macrophages. (**H**) UMAP visualization of Mo/Mac subclusters annotated by key functional genes, including *CCL2, TGFB1, IL1B, IL10, VEGFB, VEGFA, CD40*, and *CD80*. (**I**) Representative scheme and confocal images of neutrophils (in blue) injected in the vasculature (in orange) within the OrganiX^TM^. Insets highlight neutrophil localization within or adjacent to cancer cells (in green) which intravasated into the vasculature. Numbers indicate sequence of images taken at subsequent time points from Supplementary video 1. White circles: area of neutrophil-cancer cell interaction. Red arrowheads: location of first neutrophil-cancer cell interaction. Scale bar: 200µm. Schematic created in BioRender.com.

To dissect PSC responses to increasing heterogeneity, we performed GSEA on the 20 most upregulated genes of CSEM PSCs versus CS PSCs, identifying EMT, hypoxia, TNF signaling via NF-kB and glycolysis pathways (Figure 5B), similarly to signatures observed in CSEM epithelial cells. Among the most upregulated genes in CSEM PSCs compared to CS PSCs (Supplementary data file S4), we identified *WNT5A*, [38] as well as ECM-related *POSTN*, *DCN*, and *MMP9*, all linked to immunosuppression, chemoresistance, desmoplasia and cancer cell invasiveness in PDAC [39,40]. By contrast, PSCs in CS and CSE spheroids showed minimal transcriptional differences (Supplementary data file S4). These data indicate that PSCs sense the increased hypoxia of CSEM spheroids and activate WNT, metabolic and ECM-remodeling pathways associated with PDAC aggressiveness, similarly to epithelial cells. Unsupervised re-clustering of PSCs identified 4 subclusters similarly populated in the CS, CSE and CSEM spheroids (Supplementary figure S3). Pathway enrichment of upregulated genes (max 200 genes, Log_2_FC > 0.5, adjusted p value > 0.05) showed transcriptional signatures of WNT activation, ECM organization, and proliferation (Supplementary Figure S3), leading to label the subclusters as WNT-activated, Pro-fibrotic, ECM remodeling and Proliferative. WNT-activated and ECM remodeling PSCs were enriched for several collagen genes (*COL4A5, COL7A1, COL4A6, COL12A1, COL5A1*) and ECM modifiers (*LOXL2, MMP11*) (Supplementary data file S4). By contrast, the pro-fibrotic cluster showed highly expressed *TIMP1* and *TPM2* (Supplementary data file S4), consistent with a myCAF-like, contractile phenotype in PDAC [41], and *TIMP1* high expression in advanced PDAC stages and poor prognosis [42]. These PSC states underlined dual roles in the generation and preservation of deposited ECM in our spheroids, in line with their contribution to PDAC desmoplasia [43].

To align PSC phenotypes with CAF states in patient tumors we performed a supervised dimension reduction analysis using genes defining myofibroblastic CAFs (myCAFs), inflammatory CAFs (iCAFs), and antigen presenting CAFs (apCAFs) (Supplementary table S1) [44]. This analysis identified 3 subclusters (Figure 5C), with all 3 present only in CSEM spheroids and an expanded iCAFs fraction in CSEM (21.8%) compared to CS (15.3%) and CSE spheroids (16.3%) (Figure 5D), implicating monocytes/macrophages in promoting apCAF and iCAF differentiation. Spatial gene-expression profiling of vascularized CSEM spheroids further revealed CAF program heterogeneity. Whereas iCAF and classical myCAF signatures broadly distributed, CDCP1-FTL^+^ and proliferating CAF programs were enriched in proximity of the spheroids (Fibro2-dominant regions) (Figure 5E). The hypoxia-induced CDCP1-FTL CAF transcriptional program [45] presented the highest enrichment among the non-epithelial cells of the vascularized model, indicating that fibroblast states in proximity to CSEM spheroids are modulated by tumor-derived hypoxic cues.

### Heterocellular spheroids generate heterogeneous TAM populations with protumorigenic, immunosuppressive, and IL1B inflammatory phenotypes

In CSEM spheroids, monocyte derived macrophages (Mo/Mac) uniformly expressed *CD68* together with M2-associated markers *CD163* and *CD206*, indicating polarization toward an immunosuppressive TAM-like phenotype (Figure 5F). Their transcriptional profile closely matched an M2 macrophage infiltration signature from PDAC cohorts (Figure 5G), and included genes linked to inflammatory responses with both positive and negative regulation of immunity. Mo/Mac showed high expression of complement genes (*C1QA, C1QB, C1QC*), chemokines (*CXCL8, CXCL2, CCL18, CCL2, CCL3*) that promote neutrophil chemotaxis, and multiple immunosuppressive molecules (*NECTIN2, LGALS3/9, LILRB4, VSIG4, VSIR, PTPN6, TNFRSF14, MARCO, HAVCR2*), as well as the invasive front TAM marker *FOLR2* (Supplementary data file S1) [46], demonstrating faithful recapitulation of PDAC associated TAM features. Dimensionality reduction of the Mo/Mac compartment using canonical M1/M2 markers (Supplementary table S2) resolved 9 transcriptionally distinct subclusters (Figure 5H). The *IL1B* subcluster was the most divergent, with high *IL1B* and *TNF* expression and enrichment for genes associated with the annotation “cellular response to lipopolysaccharide”, consistent with a mixed inflammatory–immunosuppressive state (Supplementary Figure S4). Seven additional subclusters displayed predominantly immunosuppressive programs and were annotated by dominant genes (*CCL2, TGFBI, VEGFA, VEGFB, CD40, CD80, IL10*), while one cluster lacked a defining marker (“Unspecified”) (Figure 5H, Supplementary Figure S4). Co-expression of co stimulatory molecules *CD40/CD80* with *CD206/CD163* further supported hybrid activation states, positioning this model as a valuable tool for identifying immunoregulatory myeloid targets. Indeed, in vascularized CSEM spheroids “immunoregulating myeloid cell” signatures were detected both within vessels and in spheroid regions, with stronger enrichment in hypoxic zones (Supplementary Figure S4). Live imaging of perfused neutrophils showcased their dynamic interaction with cancer cells invading the vascular bed, consistent with high *CXCL8* expression in EMT cluster (Supplementary data file S1). Neutrophils accumulated around intravascular cancer cells and appeared to facilitate their dissemination from the primary spheroid (Figure 5I, Supplementary video 1).

### Heterocellular spheroids recapitulate desmoplastic, aggressive PDAC signaling, patient poor-prognosis programs, and drug resistance validated by functional assays

CellChat ligand-receptor interaction analysis revealed that by better mimicking the cellular complexity of PDAC TIME with CSEM spheroids translates into extensive ligand–receptor crosstalk among cancer cells, PSCs, ECs and Mo/Mac, with particularly strong communication between cancer cells and ECs mediated by *MDK, SEMA3A, NAMPT, ANGPTL4* and *VEGF* (Supplementary figure S5), all signals linked to inflammation, EMT, invasion, angiogenesis and gemcitabine resistance [29,47–49]. PSCs emerged as major sources of protumor factors, including *AREG* (EGFR-driven EMT [50] and poor prognosis [51]), *POSTN* and *WNT5A* (tumor growth, invasion, angiogenesis [49,52]). *ANGPTL4* (gemcitabine resistance [49]) engaged PSCs via *CDH11*, ECs via *CDH5* or integrins, and Mo/Mac via *SDC2* (Supplementary figure S5). Additional PSC–Mo/Mac circuits involved *MIF, MDK* and *NAMPT* (Supplementary figure S5), which drive EMT, proliferation, angiogenesis and immunosuppression [19,48,53], and complement mediated interactions between EMT cells and Mo/Mac via *SRGN* supported angiogenesis and metastasis [54]. Together, these interactions position CSEM spheroids as a desmoplastic, angiogenic, immunosuppressive and drug resistant PDAC TIME model well suited for studying cell–cell communication.

To benchmark against human disease, we compared spheroid transcriptional profiles with PDAC single-cell datasets from the Deeply Integrated human Single-Cell Omics (DISCO) platform [55], stratified by TNM stage and histological grade. Indeed, CSEM spheroids better recapitulated gene expression patterns of T3N2M0 tumors and poorly differentiated PDAC (Figure 6A). Spearman correlations showed that CSEM cells matched patient ductal, macrophage and fibroblast populations (Figure 6B). PSCs aligned with CXCL⁺, LUCAT1⁺, complement factor⁺ and collagen⁺ fibroblast subsets that sustain immunosuppression and desmoplasia [56,57], while ∼20% of spheroid macrophages mapped to SPP1⁺ TAMs associated with protumor activity and poor prognosis [58]. Thus, CSEM spheroids capture key TIME cell states observed in PDAC patients.

**Figure 6.**
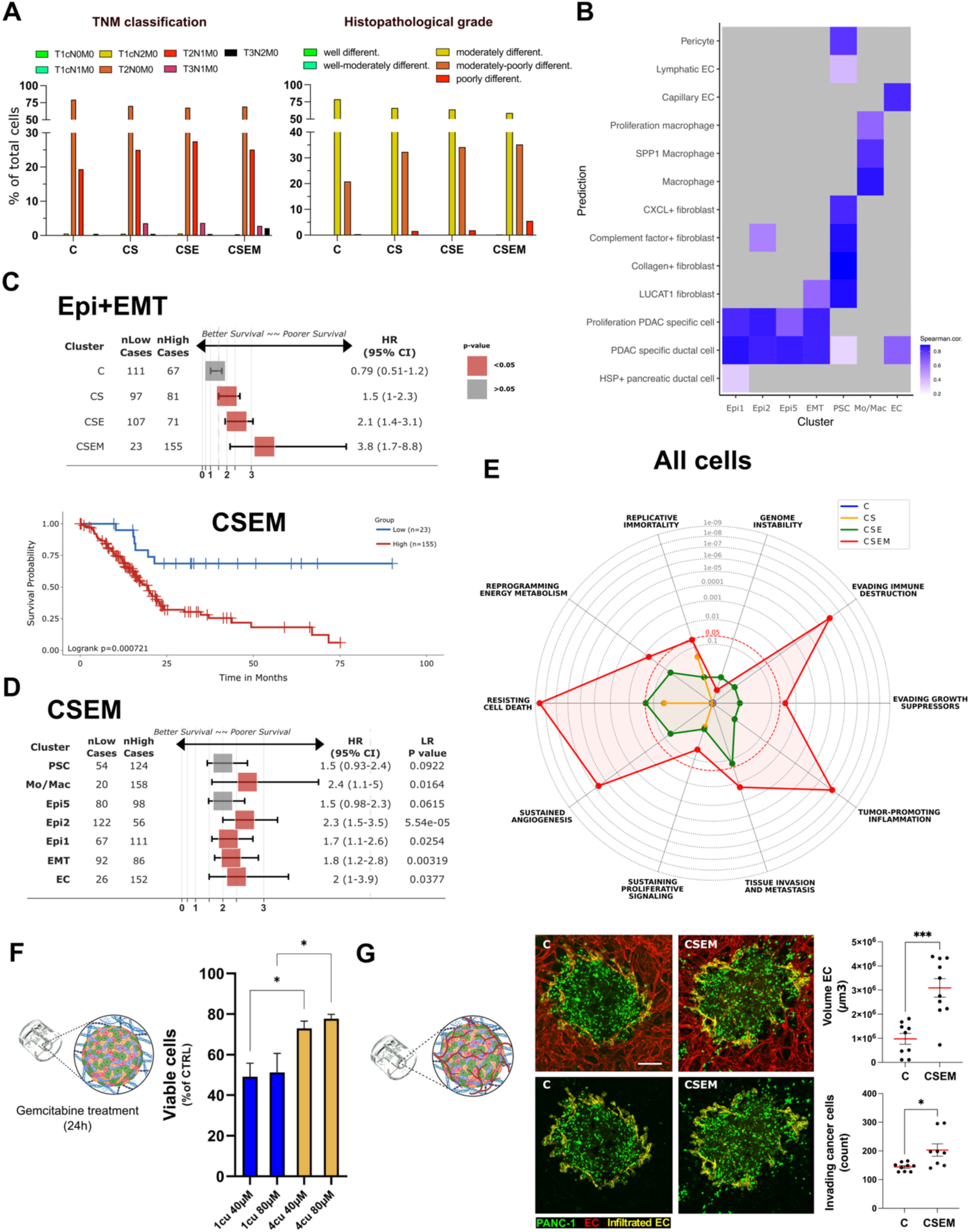
Comparison of transcriptional profiles of spheroids with PDAC patient data and their gemcitabine sensitivity. (**A**) Bar plots showing epithelial cells classified based on their resemblance to transcriptional programs. (**B**) Spearman correlation coefficients between cluster-wise expression profiles (x-axis) and curated cell-type/function gene signatures (y-axis), including pericyte, endothelial cells (lymphatic/capillary), macrophage subtypes (SPP1 and proliferative), fibroblast programs (collagen, CXCL, complement factor, LUCAT), and PDAC epithelial programs (ductal; proliferative). Color intensity reflects correlation strength. (**C**) Forest plot of the hazard ratio (HR) for the upregulated genes of the epithelial cells of the spheroids at different cellular heterogeneity (up). Kaplan-Meier plot showing survival of patients with increased expression of genes upregulated in epithelial cells of CSEM spheroids when compared to other spheroids with different composition. (**D**) Forest plot of the hazard ratio (HR) for the unregulated genes of the cell clusters identified in the CSEM spheroids. **(E)** Radar chart of the over-representation analyses (ORA) of upregulated genes in cells of spheroids at different composition. (**F**) Gemcitabine sensitivity assay on CSEM spheroids compared to C spheroid cultured in 3D microfluidic OrganiX^TM^ inserts and treated for 24h. (**G**) Representative confocal images of vascularized spheroids in OrganiX^TM^ (day 7) with quantification of volume of infiltrating ECs (yellow) and cancer cell (green) invasion. Data are shown as mean ± SEM. Statistical analysis by unpaired t test * p-value < 0.05, *** p-value < 0.001. Scale bar: 200µm. Anti-CD31 antibody (red). Schematic in F and G created in BioRender.com.

We next assessed prognostic relevance by deriving DEGs per spheroid type (all cells combined) and computing survival associations using GSVA based high versus low enrichment scores. The CSEM signature was linked to shorter survival with the strongest impact (HR 8.8), at least 6- and 4-fold higher than CS and CSE, respectively (Supplementary Figure S6). Restricting to epithelial DEGs, cancer cells from CSEM spheroids remained the most correlated to poor prognosis (HR 3.8 vs 2.1 for CS and 1.5 for CSE) (Figure 6C), as also illustrated by Kaplan–Meier curves distinctly separating patients with high versus low CSEM epithelial signatures, with patients harboring high CSEM signatures exhibiting the lowest survival probability. Cluster level GSVA showed that Mo/Mac and Epi2 contributed most to the poor survival signature, followed by EC, EMT and Epi1 (Figure 6D), indicating that increasing cellular complexity drives a transcriptional landscape in CSEM spheroids that most closely mirrors poor-prognosis PDAC. Over representation analysis (ORA) of genes upregulated across whole CSEM spheroids identified 7 cancer hallmarks uniquely enriched in this model (resistance to cell death, tumor promoting inflammation, evasion of growth suppressors and immune destruction, invasion/metastasis, angiogenesis and metabolic reprogramming), consistent with an aggressive PDAC TIME (Figure 6E). Functionally, CSEM spheroids were more chemoresistant to 24 h gemcitabine treatment, with a significantly higher fraction of viable GFP–PANC 1 cells in CSEM than in C spheroids (Figure 6F). To functionally validate the angiogenic and invasive programs identified molecularly, we vascularized C and CSEM spheroids in OrganiX™ microfluidic chips, generating perfusable endothelial networks that supported PBMC or neutrophil flow for 24 h (Supplementary video 1, Figure 6G). In CSEM spheroids, vessels penetrated the tumor mass, whereas in C spheroids ECs remained largely peripheral; volumetric analysis confirmed significantly greater intratumoral vascular volume in CSEM spheroids (Figure 6G, upper panels). Tracking GFP–PANC-1 dissemination revealed that CSEM spheroids released more invading cancer cells from the primary mass than C spheroids, consistent with higher metastatic potential (Figure 6G, lower panels). Collectively, these data show that CSEM heterocellular spheroids integrate disease relevant signaling circuits, poor prognosis transcriptional programs and reduced drug sensitivity, providing a robust preclinical model for aggressive PDAC.

## Discussion

Our heterocellular PDAC spheroid model integrating cancer cells, PSCs, Mo/Mac, and ECs into a vascularized microphysiological system, recapitulates key transcriptional programs of aggressive, poor-prognosis PDAC. The platform embeds multiple cell types providing a more realistic TIME, and, through vascularization, enables the study of leukocyte recruitment and interactions (neutrophils, PBMCs). The scRNA-seq and spatial transcriptomics suggest hypoxia as a central driver of CSEM spheroid features, mirroring the aggressive and chemoresistant PDAC hallmarks. Hypoxia contributes to inflammation, EMT, and metabolic changes through stromal-immune crosstalk [36,59]. The differential gene expression analysis of the Epi1 cluster (scRNA-seq) and CC1 (spatial analysis) in CSEM spheroids shows a higher expression of hypoxia-related genes like *HIF1A* and *ANGPTL4*, and upregulated glycolysis-related genes, such as *LDHA*, and *PGK1*. Proliferative Epi2 cancer cells (scRNA-seq) and CC5 (spatial analysis) cluster populated the area with stronger enrichment for proliferation-related genes (G2M checkpoint gene set), locating at the periphery of the CSEM spheroids, adjacent to the hypoxic CC1 cluster, and in small aggregate which have departed from the parent spheroid. These findings align with patient-derived PDAC xenograft models, in which hypoxic regions co-localize with lactate accumulation, promoting the proliferation of normoxic neighboring cells [60]. Accordingly, CSEM spheroids appear to recapitulate spatial cell proximity-dependent effects, whereby hypoxia-associated metabolites influence neighboring cells and affect key transcriptional programs toward enhanced tumor growth. The higher percentage of Ki67^+^ cancer cells observed by immunostaining in our CSEM spheroids [13], provides further evidence of the contribution of heterogeneity in modulating cancer cell proliferation. CellChat analysis of our CSEM spheroids predicts multiple ligand–receptor interactions consistent with downstream activation of hypoxia-responsive transcriptional programs. CSEM cancer cells presented high *ANGPTL4* expression, a hypoxia (via HIF-1α) inducible gene in pancreatic cancer cells [61]. *ANGPTL4* expression has been associated with aggressive basal/squamous-like tumor cell phenotype, poorer survival in TCGA PDAC RNA-seq cohorts, and gemcitabine chemoresistance [49,62], consistent with our functional assays results showing gemcitabine resistance and higher cancer cell invasion in CSEM spheroids. In line with these observations, patients with high CSEM epithelial signatures exhibited lower overall survival. Other strong ligand-receptor interactions suggestive of a hypoxic-driven aggressive basal/squamous cell phenotype involved *SEMA3A, NAMPT* and *VEGF*. Indeed, in the vascularized CSEM model most of the cancer cells present a basaloid signature, such as the one observed in patients with poor prognosis, and ligand-receptor interaction analysis suggests stromal-immune crosstalk. Notably, the Fibro2 cluster potentially amplifies hypoxia-induced ANGPTL4 via PGE₂, analogous to observations in colorectal cancer [63].

Additional cellular interactions identified from the incoming and outgoing signaling pattern analysis strengthen our molecular findings about inflammation, metabolic alterations, metastasis, angiogenesis, and therapy resistance. The CSEM spheroid environment was characterized by ligand-receptor interactions of inflammatory molecules, such as *SRGN* [54], *RARRES2* (chemerin) [64], *IL-16* [65], and *NAMPT [48]*, and stroma-tumor interactions promoting metastasis via *AREG* and *ANXA1 [50,66]*. The differential regulation of these genes has been demonstrated to be a downstream effect of hypoxia, orchestrating different protumor activities that enhancing chemoresistance, tumor cell stemness, metastasis, angiogenesis, and fibroblast differentiation in PDAC or other cancers [26,54]. These findings reinforce our hypothesis that hypoxia orchestrates cell-cell interactions determining a more aggressive TIME. As mentioned, the interplay among different cell types in the TIME of the CSEM spheroids well replicates glycolytic signatures of cancer cells in PDAC. Specifically, the *PGK1* upregulation suggests intercellular signaling pathways between Mo/Mac and other cell populations, driving glycolysis through NF-kB signaling. In PDAC, monocyte-derived CCL18 was identified as an upstream activator of NF-kB in cancer cells expressing vascular cell adhesion molecule 1 (VCAM1), increasing the interaction with monocytic cells and promoting the Warburg effect [67], in agreement with evidence that tumor-conditioned macrophages adopt a Warburg-like metabolic program that enhances metastasis in PDAC [68]. Consistently, the high expression of *CCL18*, a M2 macrophage marker [69], by Mo/Mac (∼28 fold higher than other cell clusters) provides a reasonable explanation for their role in enhancing the NF-kB-activation and promoting glycolysis in the CSEM model. Our previous study confirms that Mo/Mac presence in the spheroid is a requisite for the increased concentration of IL-6 in the supernatant, a clear downstream effect of NF-kB activation [13]. Macrophage expression of CCL18 requires oxygen [69], and accordingly, we identified *CCL18*+ myeloid cells mainly at the periphery of the spheroids (Supplementary figure S7) but not in the core. Therefore, our model recapitulates CCL18-driven aggressive features of PDAC. In line with this, NF-kB pathway activation was observed in cancer cells, PSCs, and Mo/Mac, and this was greatly enhanced in CSEM spheroids compared to lower-heterogeneity models, confirming Mo/Mac as pivotal players in establishing an inflammatory environment and contributing to metabolic reprogramming in PDAC [67]. Proteomic profiling provided validation evidence supporting the transcriptional and functional data, further positioning the CSEM system as a model of aggressive PDAC. The upregulated proteins observed in CSEM cancer cells, were consistent with a highly proliferative state coupled to an enhanced autophagic program (ATG7 and SQSTM1), this latter activated by hypoxia and ER stress among other cell stresses [70]. We also detected increased major histocompatibility complex class I (MHCI) antigen-processing components (TAP1/2, TAPBP), suggesting upregulated machinery for MHCI/peptide assembly, downstream of IFN signaling in our immunocompetent CSEM spheroids. This evidence is in line with *in vivo/in vitro* PDAC CRISPR screens showing TAP1/TAPBP/ATG7 as immune-context dependencies that are diminished/dispensable *in vitro*, consistent with these programs being induced by immune-derived IFN cues [71]. Functionally, *ATG7*-linked stress tolerance and anti-apoptotic wiring (especially in Epi1 and EMT clusters of CSEM) represents a combination that can support survival under cytokine/TNF pressure and provides a plausible explanation for why higher *TAP1* associates with poor prognosis and therapy resistance [72,73], again highlighting CSEM model molecular similarity to PDAC.

Interestingly, our highly invading CSEM spheroids showed that EMT (scRNA-seq) and CC5 (spatial analysis) cluster gene signatures suggest a role for these cells in establishing an inflammatory TIME. Indeed, the CSEM invading cells (CC5/EMT) activated pathways associated with TNF-mediated NF-kB translocation and necroptosis, likely contributing to the inflammatory microenvironment that promotes tumor invasiveness and an immunosuppressive milieu in PDAC [74]. In line with their spatial distribution, EMT cells upregulated the expression of invasion markers, such as *IL-32, SDC4* and *LCN2* [25–27], reinforcing the hypothesis that our CSEM model recapitulates, at the molecular level, features of invasive PDAC as a downstream effect of the hypoxia-glycolytic-mediated inflammation. The increased invasiveness of our CSEM spheroids was well supported by the similarities between the gene expression profiles of patient-derived invasive cells and the CSEM EMT cells, including the p53 pathway activation. Consistently, our immunofluorescence images showed that the invasive side of CSEM spheroids was strongly expressing p53 (mutant in PANC-1 cells) which has been demonstrated to promote invasiveness [75]. In contrast, p53 was not expressed in the core of CSEM spheroids, in line with previous findings showing p53 mRNA downregulation in hypoxic PANC-1 cells [76]. Notably, the EMT cells resemble hybrid cells expressing both epithelial and leukocyte markers, such as *KRTs* and *CD45*, suggesting potential fusion mechanisms between Mo/Mac and cancer cells which are known to enhance metastatic potential and immune evasion [28].

Our CSEM PSCs also recapitulate the transcriptional signature of desmoplastic PDAC by upregulation of ECM-related genes such as *DCN* [40] and *POSTN* [39], known to confer aggressive features to cancer cells [40] and immunosuppression [39]. Spatial analysis identified a higher degree of similarities between our fibroblastic cells and the hypoxia-induced CDCP1-FTL^+^ fibroblasts observed in patients, highlighting intercellular communication affecting distal cells around the hypoxic spheroid. The presence of these hypoxia-induced fibroblasts correlates with poor clinical outcome in PDAC patients, suggesting that the vascularized model mimics features of aggressive and chemoresistant pancreatic cancer. Consistently, CSEM spheroids have an increased transcriptional similarity with cells of poorly differentiated tumors and more advanced PDAC TNM stages. The comparison between the gene expression profiles of each primary cell cluster of our CSEM spheroids and patient-derived cells highlighted similarities between their expression patterns and key progression-associated cells. Strikingly, the human primary cells (i.e., PSCs and monocytes) used to form our CSEM spheroids adapted their gene expression to the tumor environment to transcriptionally resemble different clusters of cells from the patient primary tumor rather than adjacent normal tissue (Supplementary figure S5). Notably, specific protumor and pathologically relevant clusters, such as SPP1^+^ macrophages, complement secreting LUCAT1^+^ fibroblasts [56,57], were partially recapitulated by the primary cells embedded in our CSEM spheroids. This well corroborate with the CSEM cells upregulating genes which in patients associate with a decrease in their overall survival. Moreover, the ORA using the upregulated genes of our spheroids with different cellular heterogeneity confirmed that our CSEM model enhanced the protumor functions associated with poor prognosis in patients. Taken together, these results clearly encourage the use of this system for the identification of microenvironmental targets to counteract tumor promoting cellular functions.

The lower sensitivity of CSEM spheroids to gemcitabine treatment compared to monoculture spheroids confirmed that higher-heterogeneity spheroids better mimic the chemoresistance that is often observed in patients and leads to poor prognosis. Similarly to the resistance to cell death, the other cancer hallmarks found enhanced in CSEM spheroids through the ORA analysis, namely tissue invasion and sustained angiogenesis, were confirmed by our functional assays. CSEM spheroids showed increased infiltration of ECs in the tumor aggregate and cancer cell migration, making this model a powerful platform for linking molecular and functional findings. Invasive EMT cells in CSEM spheroids express high levels of *CXCL8* and *CXCL2*, both neutrophil-recruiting cytokines. Indeed, when we challenged the vascularized model with neutrophils, within the vascular bed we observed physical cellular interactions between neutrophils and the cancer cells migrating from our CSEM spheroids, recapitulating the tumor-infiltrating interactions between neutrophils and invading cancer cells *in vivo* [77,78] . This interaction between evading cancer cells and neutrophils has been inferred in several studies [77,78] but in this study it is directly visualized in a complex human 3D vascularized *in vitro* model resembling key aspects of PDAC malignancy. The model enables unprecedented visualization of neutrophil-cancer cell dynamics, which can now be studied molecularly in a relevant human PDAC immune microenvironment.

## Conclusion

The vascularized PDAC heterocellular spheroid model presented in this study recreates the PDAC niche *in vitro* using non–patient-derived cells, offering a scalable and tunable alternative for mechanistic and translational studies. Patient-derived components, although physiologically relevant, often are limited and lack space for scalability, reproducibility, and systematic dissection of individual cell-type contributions. These constraints are particularly critical when rapid prediction of patient response is needed within a clinically relevant timeframe, or when larger cohorts are required for robust mechanistic hypotheses. In contrast, our model recreates the essential features of the PDAC niche entirely *in vitro* using non–patient-derived cancer, stromal, endothelial, and immune cell sources. This approach enables scalable, reproducible generation of complex spheroids with controllable cellular composition, overcoming the bottlenecks of patient tissue dependence. By combining single-cell transcriptomics, spatial biology, proteomics, and microfluidic-based functional assays, our platform provides insights on cell–cell signaling, spatial organization, and molecular adaptation. This integrative, multi-omics strategy sheds new light on how specific cell populations orchestrate tumor behavior and therapy resistance, offering a robust, tunable alternative for mechanistic investigation and translational drug development in PDAC research.

## Materials and Methods

### Cell culture

PANC-1 (American Type Culture Collection, ATCC) were maintained in Iscove Modified Dulbecco Media (IMDM) supplemented with 10% fetal bovine serum (FBS, Thermo Fisher Scientific, Waltham, MA, USA, Cat. no. 10082147) and penicillin/streptomycin (100 U/mL, Invitrogen/Gibco Cat. no. 15140122). Human Pancreatic Stellate Cells (HPaSteC, here referred as PSCs) (Gene Etichs Cat. no. 3830, Lot. no.14358) were maintained in Stellate Cell Medium (SteCM, ScienCell Research Laboratories, Carlsbad, CA, USA, Cat. no. 5301) supplemented with 2% FBS (ScienCell Research Laboratories, Cat. no. 0010), 1% stellate cell growth supplement (ScienCell Research Laboratories, SteCGS, Cat. no. 5352) and 1% antibiotic solution (ScienCell Research Laboratories, Cat. no. 0503). Human umbilical vein endothelial cells (HUVECs, here referred as ECs) (Lonza, Basel, Switzerland, C2519AS, Lot. no.633426; Promocell, Heidelberg, Germany, Cat. no. C-12205) were cultured in EGM-2™ SingleQuot™ containing 0.5 ng/ml VEGF, 5 ng/ml EGF, 10 ng/ml bFGF, 20 ng/ml long R3-IGF-1, 22.5 mg/ml heparin, 1 mg/ml ascorbic acid, 0.2 mg/ml hydrocortisone, gentamicin (1/1000 dilution) and 2% FBS. For vascularization cells were cultured in the OrganiX^TM^ insert (AIM-Biotech, Singapore) with EGM-2^TM^ MV Microvascular Endothelial Cell Growth Medium-2 BulletKit^TM^. Normal human lung fibroblasts (NHLF, Lonza Cat. no. CC-2512. Lot. no. 21TL019232) were maintained in Dulbecco’s Modified Eagle Medium (DMEM) supplemented with 10% fetal bovine serum (FBS, Thermo Fisher Scientific Cat. no. 10082147), penicillin/streptomycin (100 U/mL, Invitrogen/Gibco Cat. no. 15140122), Glutamax (dilution 100x, Invitrogen/Gibco, Thermo Fisher Scientific, Cat. no. 35050061) and sodium pyruvate (1 mM, Invitrogen/Gibco, Cat. no. 11360070). All cells were cultured in 75 and 175 cm2 tissue culture treated flasks in a humidified atmosphere composed of95% air and 5% CO2 and a temperature of 37°C. Cells were passaged every 72 h using 0.25% (PANC-1 and HPNE) or 0.05% (ECs and PSCs) Trypsin- EDTA (Gibco, Thermo Fisher Scientific, Cat. no. 25300054).

### PBMC, neutrophils and monocyte isolation for spheroid formation and functional assays

Peripheral blood mononuclear cells (PBMC) and neutrophils were isolated from blood cones (for spheroid formation) or from healthy donor blood collected in sodium citrate tubes (0.109 M, 3.2%, BD, Franklin lakes, NJ, USA, BD Vacutainer, Cat. no. 363083) for functional assays. De-identified human blood tissue was collected from blood cones in accordance with and under the following project: HSA Residual Blood Samples for Research, project titled “Harnessing immune response for new therapies in transplantation and cancer” (A*STAR IRB Reference Number: 2024-133). De-identified human blood tissue was collected from healthy donors in accordance with the approved IRB studies titled “Study of blood cell subsets and their products in models of infection, inflammation and immune regulation” (CIRB Ref: 2017/2806 and A*STAR IRB Ref. No. 2024-003). We used a density gradient of Ficoll/Paque PLUS (GE Healthcare, Marlborough, MA, USA) centrifuging at 900 x g, at room temperature (RT) for 20 min. The obtained PBMC layer was collected using a sterile pipette and washed with Ca^-^Mg^-^ PBS before incubation with red blood cell lysis (155 mM NH_4_Cl, 10 mM KHCO_3_, 0.1 mM EDTA) for 5 min at RT. The pellet below the PBMC layer was diluted with red blood cell lysis and kept on ice for 20 min. At the end of the respective incubations, cells were washed in Ca^-^ Mg^-^ PBS. PBMC from blood cones were prepared for cryopreservation using Bambanker™ (Fujifilm Wako Chemicals U.S.A. Corporation, Richmond, VA, USA). Neutrophils and PBMC from blood tubes were counted and resuspended in EBM2 containing 5% pooled human serum (Sigma-Aldrich, MO, USA, Cat. no. H5667, Lot. No. SLCF0860) and prepared for the functional assays at a concentration of 4.3 x 10^6^ cells/mL. On the day of the experiment, cryopreserved PBMC suspension was used for the isolation of monocytes using Pan Monocyte Isolation kit (Miltenyi Biotech, Bergisch Gladbach, Germany). The isolated cells were characterized by the expression of CD14 and CD16 in a CD45+/CD3- gate. Monocytes represented more than 95% of the total CD45^+^ cells.

### Single-cell RNA sequencing

Multiple spheroids (60 for each condition) were collected in an Eppendorf tube 7 days after cell seeding in hanging drops (Figure 1). Spheroids were incubated with collagenase IV (D) in Roswell Park Memorial Institute (RPMI, serum free) for 6 min at 37⁰C. Trypsine/EDTA solution (0.25%) was added in an equal volume and incubated for additional 5 min resuspending (at least once) using a 200µl pipette tip during the incubation. After 5 min EGM2 was added to stop the enzymatic digestion and cells washed once with PBS + 1% BSA. Single-cell suspensions of spheroid-derived cells in PBS + 1% BSA were loaded for droplet encapsulation using the 10x Genomics Chromium Controller at a targeted cell recovery of about 6,000 cells per sample. Single-cell gene expression libraries were prepared using the 10x Genomics Chromium Next GEM Single-Cell 3′ v3.1 Library & Gel Bead Kit according to the manufacturer’s protocol. The libraries were subjected to 2 × 151 cycle sequencing on an Illumina Novaseq 6000 at a targeted sequencing depth of 50,000 reads/cell.

### Single-cell mRNA mapping

Cell Ranger (version 7.0.0) was used to map single-cell RNA-sequencing reads of each sample to the human reference genome version GRCh38 release 93 (GRCh38-3.0.0 reference package downloaded from 10x Genomics). All samples passed the minimum quality settings.

### Seurat analysis

The raw feature count matrices were imported and analysed using the Seurat package (version 4.1.1) [79] on the R platform (version 4.2.0, http://www.r-project.org/). All samples were filtered to retain cells with more than 500 RNA features and less than 15% mitochondrial RNA content. For each sample, we applied the standard Seurat processing pipeline of log normalization with the “NormalizeData” function (“LogNormalize” option and scale factor of 10,000), variable feature selection with the “FindVariableFeatures” function (“vst” as the method and 2,000 as the number of features selected), and data scaling with the “ScaleData” function applied to all genes). Thereafter, we performed dimension with PCA and UMAP with the “RunPCA” and “RunUMAP” functions, respectively. As inspection of the UMAP did not suggest batch effects present, no data integration was performed. We then merged all samples into one Seurat object and again applied the standard Seurat processing pipeline. To obtain clusters, we employed the “FindNeighbors” followed by “FindClusters” functions. At a resolution of 0.1, we obtained seven clusters. For annotation, DEGs were computed using the “FindMarkers” and “FindAllMarkers” functions for pairwise comparisons and one sample vs the rest, respectively. Using known markers, we annotated three epithelial clusters, one cluster each of mesenchymal, fibroblast, macrophage, and endothelial cells. We then employed the same Seurat processing pipeline to each cell type-specific clusters for additional analysis.

### Cell-cell communication prediction with CellChat

We employed the CellChat R package (version 1.6.1) to predict cell-cell communications, following the accompanying tutorial and using the default parameters.

### Sample preparation for proteomic analysis

Multiple spheroids (300 for each condition) were collected in an Eppendorf tube 7 days after seeding in the hanging drop device (Figure 1). Spheroids were incubated with collagenase IV (D) in Roswell Park Memorial Institute (RPMI, serum free) for 6 min at 37 ⁰C. Trypsin/EDTA solution (0.25%) was added in an equal volume and incubated for additional 5 min resuspending (at least once) using a 200 µl pipette tip during the incubation. After 5 min EGM2 was added to stop the enzymatic digestion and cells washed with PBS + 1% BSA. DAPI was added to the cell suspensions before fluorescence-activated cell sorting to exclude dead cells. Enough PANC-1 cells were sorted using their intrinsic expression of GFP PANC-1 cells were lysed with 8M urea, 50 mM ammonium bicarbonate (pH 8, Sigma-Aldrich), 0.5% sodium deoxycholate, supplemented with 50 µg/mL DNase I (Sigma-Aldrich) and RNase I (Thermo Fischer Scientific), EDTA-free protease and phosphatase inhibitors (cOmplete™, Mini, Roche Diagnostics). Lysates were cleared by centrifugation at 21,300g for 1 hour at 16°C. Protein content of the lysed samples was determined using Bradford protein assay (Bio-Rad, USA). Samples were reduced with 10 mM dithiothreitol (DTT) at room temperature for 60 min, and alkylated in the dark with 20 mM iodoacetamide (IAA) at 20 °C for 30 min. An additional final concentration of 10 mM DTT was added to quench the excess IAA. The samples were diluted with 50 µM ammonium bicarbonate to achieve a final concentration of 2 M urea; followed by digestion with trypsin (Sigma; 1:30 enzyme-to-protein ratio) at 37 °C, overnight. The digestion was quenched with 1% formic acid, digested peptides were desalted by Sep-Pak C18 1 cc Vac cartridges (Waters, USA). Detergent in the sample was cleaned by cation exchange using SCX SPE Tube (Supel™, Select, Supelco, Bellefonte, PA, USA). The samples were dried and stored at -80 °C for further LC-MS/MS analyses.

### LC–MS/MS analyses

Peptides were reconstituted in 2% acetonitrile, 0.06% trifluoroacetic acid and 0.5% acetic acid. The measurements were performed in data-dependent acquisition mode on an Orbitrap Eclipse Tribrid Mass Spectrometer (Thermo Fisher Scientific) coupled with a Vanquish™ Neo UHPLC System (Thermo Fisher Scientific). Solvents used were 0.1% formic acid in water (Solvent A) and 0.1% formic acid in 80% acetonitrile (Solvent B). Samples were analyzed in quadruplicates with a 180-min gradient of Solvent B on an analytical column (2 µm C18, 75 µm X 500 mm, Thermo Fisher Scientific). More details can be found in the extended methods section in supplementary material.

### Stereo-sequencing (stereo-seq)

Stereo-seq experiment, library preparation, and sequencing were performed as described in an established pipeline [80]. OCT compound-embedded tissue blocks were cut into 15 μm thick sections. Tissue permeabilization was tested with varying durations (3, 6, 12, 18, 24 min) on the Stereo-seq Chip P Slide using the 15 µm tissue sections. The optimal permeation time was determined to be 24 min. Following cryosection, tissue sections are mounted on Stereo-seq chips, incubated at 37 °C, and fixed with -20 °C methanol. The chips underwent ssDNA staining and imaging with an Olympus IX83 Inverted Slide Scanning Microscope. Tissue permeabilization was performed with 0.1% pepsin in HCl buffer, followed by washing with SSC buffer containing RNAse inhibitor. Captured mRNA was reverse transcribed at 42 °C. Post in situ reverse transcription, the chips were treated with tissue removal buffer and then washed. cDNA was released from the chips using a cDNA Release Mix at 55 °C, followed by fragmentation, amplification, and purification to produce a DNA Nanoballs (DNBs) library. The DNBs were sequenced on the MGI DNBSEQ-T1 platform.

### Stereo-seq raw data processing (Alignment and UMI counting)

The Stereo-seq FASTQ data contains unique identifiers unique molecular identifier (UMI) (25 bp CID and 10 bp MID) in the first read and cDNA sequences in the second read. We used the SAW workflow (https://github.com/BGIResearch/SAW) to process this data, which involves several key steps, including mRNA spatial location restoration, filtering, mRNA genome alignment, gene region annotation, MID (Molecule Identity) correction, expression matrix generation, and tissue region extraction. The raw reads fastq files from Streo-seq were processed following the STOmics SAW (https://github.com/STOmics/SAW) pipeline. SAW count pipeline outputs gene expression matrix data at various bin and single-cell segmented level in .gef and hd5 file format. The adjusted.cellbin.gef file was subsequently used for analysis using R programming environment. More details can be found in the extended methods section in supplementary material.

### Spheroid embedding in OrganiX without vasculature

Spheroids were collected 7 days after seeding in the hanging drop device and singularly allocated to 200 µL Eppendorf and prepared for vascularization. Resuspension medium was prepared using EGM2 MV solution supplemented with 0.8% rat tail collagen type I (Corning, NY, USA, 3.41 mg/mL, Cat. no. 354236), 1 U/mL thrombin (from bovine plasma, Sigma-Merck, Cat. no. T9549-100UN). After washing with PBS, each spheroid was mixed with 25 µl of resuspension medium and an equal volume of 6 mg/ml fibrinogen in PBS. After mixing by resuspension, the spheroid suspension mix was quickly injected into the OrganiX^TM^ gel inlet and allowed to polymerize for 20 min at room temperature. After polymerization, 35µL of EGM-2 MV was added to each microfluidic device medium inlets adjacent to the gel containing the spheroid. Additional 400 and 200 µL of EGM2 MV were added to each medium chambers to create a flow within the gel. EGM2 MV+VEGF medium was refreshed daily.

### Spheroid vascularization and leukocyte injection

Spheroids were collected 7 days after seeding in the hanging drop device and singularly allocated to 200 µL Eppendorf and prepared for vascularization. Resuspension medium was prepared using EGM2 MV solution supplemented with 0.8% rat tail collagen type I (Corning, 3.41 mg/mL, Cat. no. 354236), 1 U/mL thrombin (from bovine plasma, Sigma-Merck, Cat. no. T9549-100UN) containing ECs and NHLF (ratio 4:1). After washing with PBS, each spheroid was mixed with 25µl of resuspension medium and an equal volume of 6 mg/ml fibrinogen (Type I-S: from bovine plasma, Sigma-Merck, Cat. no. F8630-5G) in PBS. The spheroid suspension mix was quickly injected into the OrganiX^TM^ gel inlet and allowed to polymerize for 20 min at room temperature. After polymerization, 35 µL of EGM-2 MV supplemented with vascular endothelial growth factor (VEGF, Recombinant human protein, Gibco, Cat. no. PHC9391) was added to each microfluidic device medium inlets adjacent to the gel containing the spheroid. Additional 400 and 200 µL of EGM2 MV+VEGF were added to each medium chambers to create a flow within the gel. EGM2 MV+VEGF medium was refreshed daily for 4 days (day 7 to day 10) before seeding ECs (1 x 10^6^/mL) in the medium inlets. After seeding, EGM-2™ SingleQuot™ was used to replenish the medium chambers with 400 and 200 µL and refreshed daily for other 2 days. On day 13, medium was changed to EBM2 supplemented with 5% human serum (type AB, Sigma-Aldrich, Cat. no. H5567-100ML). On day 14, PBMC or neutrophils were resuspended in EBM2 supplemented with 5% human serum to a concentration of 4.3 x 10^6^/mL. Thirty-five µL of the cell suspension (PBMC or neutrophils) was injected into the medium inlet. To establish a differential volume between the two chambers, 200 µL of medium was transferred from the chamber opposite to the one where the cells were injected.

### GFP-PANC-1 generation, cell labeling and immunofluorescence

GFP-tagged PANC-1 cells were generated using Lipofectamine 2000-mediated transfection following manufacturer’s instructions. Enhanced GFP (eGFP) plasmid (pEGFP-C1 EGFP-3XNLS, Cat. no. #58468, Addgene) was used for transfection. GFP-tagged PANC-1 cells were FACS sorted to enrich the positive population. For cell labeling in functional assays, cells were collected, counted, and washed once in PBS. Cells were spun down at 300 × g for 5 min and the supernatant discarded. PBS containing CellTrace™ Violet (Cat. no. C34557, Thermo Fisher Scientific, dilution 1/1000) was used to stain the different cell types. The cell pellet was resuspended in PBS containing the dyes and incubated at 37°C. After 30 min, 5 ml of medium containing human serum was used to stop the staining reaction and cells were washed before resuspending them in EBM-2 + 5% human serum for injection in OrganiX™. For immunofluorescence, non-vascularized and vascularized spheroids embedded in gel within the OrganiX™ microfluidic device were washed with PBS by introducing differential volumes into the two medium chambers, generating a flow through the gel matrix. After washing, fixation with paraformaldehyde (PFA) 4% in PBS for 20 min was performed. For extracellular staining, the devices were incubated with 5% BSA in PBS for 2 hours at room temperature. After washing with PBS, the devices were incubated overnight with the anti-human CD31 Alexa Fluor 594-conjugated antibody (Biolegend, San Diego, CA, USA, Cat. no. 102520) solution in 1% BSA PBS at 4⁰C. The antibody solution was washed away with PBS and the devices prepared for microscopy acquisition. For intracellular staining, devices were incubated using 0.5% Triton-X-100 in PBS. After 20 min, devices were washed two times with PBS before incubating them with 5% BSA in PBS for 2 hours at room temperature. Upon washing, overnight incubation with mouse anti-human p53 (DO-1) primary antibody (Santa Cruz Biotechnology, Dallas, TX, USA, Cat. no. SC-126) in 1% BSA PBS was performed at 4⁰C. Devices were washed with PBS and subsequently incubated with a blocking solution containing 3% BSA + 3% goat serum in PBS for 2 hours at room temperature. The blocking solution was washed using PBS and devices were incubated overnight with goat anti-mouse IgG (H+L) Alexa fluor 546 conjugated antibody (Invitrogen, Cat. no. A-11030) solution in 1% BSA +1% goat serum in PBS supplemented with Hoechst 33342 (1 µg/ml, Invitrogen, Cat. no. H3570). Devices were washed using PBS and prepared for microscopy.

### Microscopy

Microscopy images of the spheroids (vascularized and non) were acquired using the inverted confocal microscope Olympus FV3000. Images were processed using IMARIS software (v.9.7.1, Bitplane). We identified GFP-PANC1 using the functions “spots” selecting quality of GFP using background subtraction and considering the fluorescent signal elongation due to the acquisition. EC infiltration in the spheroids was identified using the function “surface” to create a mask for the spheroid using GFP-PANC1 signal. EC surface was created using CD31 signal. Merging volumes between the surface of the spheroids and the ones of the ECs was quantified.

### Kaplan-Meier plotter and gene hazard ratio

Differentially expressed genes (DEGs) identified between spheroids of different cellular composition were evaluated for prognostic relevance using SurvivalGenie 2.0 (https://bhasinlab.bmi.emory.edu/SurvivalGenie2), which implements publicly available code (GitHub;[81]). Overall survival (OS) in pancreatic ductal adenocarcinoma was assessed using the “Cluster Marker” module and the “TCGA-PAAD” cohort. Patients were stratified into high- and low-expression groups for each gene using the optimal cut-point option [82], and Kaplan–Meier OS curves were generated. Hazard ratios (HRs) (with confidence intervals) were obtained from Cox proportional hazards modeling and summarized as forest plots for cluster-level gene panels, with representative genes additionally shown as Kaplan–Meier curves. HR values were interpreted as the direction and magnitude of association between gene expression and OS.

### Statistics of functional assays

Data are presented as means ± standard error of the mean (SEM) if not differently stated. Statistical significance for comparisons was determined by one-way ANOVA with different post-hoc tests. A p-value less than 0.05 was considered statistically significant. All data are analyzed using GraphPad Prism (version 6.07) software (San Diego, CA, USA).

## Author contributions

GG conceived and designed the study, collected the data, directed extrapolation of data and performed PEA and GSEA, interpretation of data, performed the functional assays, and wrote the original draft. AKS and JC performed scRNAseq data extrapolation. PK performed spatial transcriptomics data extrapolation. GT and BKZX, contributed to perform experiments. CXT, EK, RB, and WW performed the proteomic analyses. AT and SWH, performed the scRNAseq. GA directed and supervised the study, and revised the original draft. WW, PC and SA revised and edited the original draft. All the authors approved the submitted manuscript.

## Ethical approvals

De-identified human blood tissue was collected from blood cones in accordance with and under the following project: HSA Residual Blood Samples for Research, project titled “Harnessing immune response for new therapies in transplantation and cancer” (A*STAR IRB Ref. No. 2024- 133). De-identified human blood tissue was collected from healthy donors in accordance with the approved IRB studies titled “Study of blood cell subsets and their products in models of infection, inflammation and immune regulation” (CIRB Ref: 2017/2806 and A*STAR IRB Ref. No. 2024-003).

## Data and materials availability

The mass spectrometry proteomics data have been deposited to the ProteomeXchange Consortium with the dataset identifier PXD058039.

