## Supplementary material for "Multi-omics and functional analysis of a bioengineered vascularized pancreatic cancer model reveal an immunosuppressive and therapy-resistant niche": Suplemental Material

**This PDF file includes:**

Figures S1 to S7
Table S1 and S2
Extended Methods

**Other supplementary materials (provided as separate files)**

Supplementary video 1
Data files S1 to S4

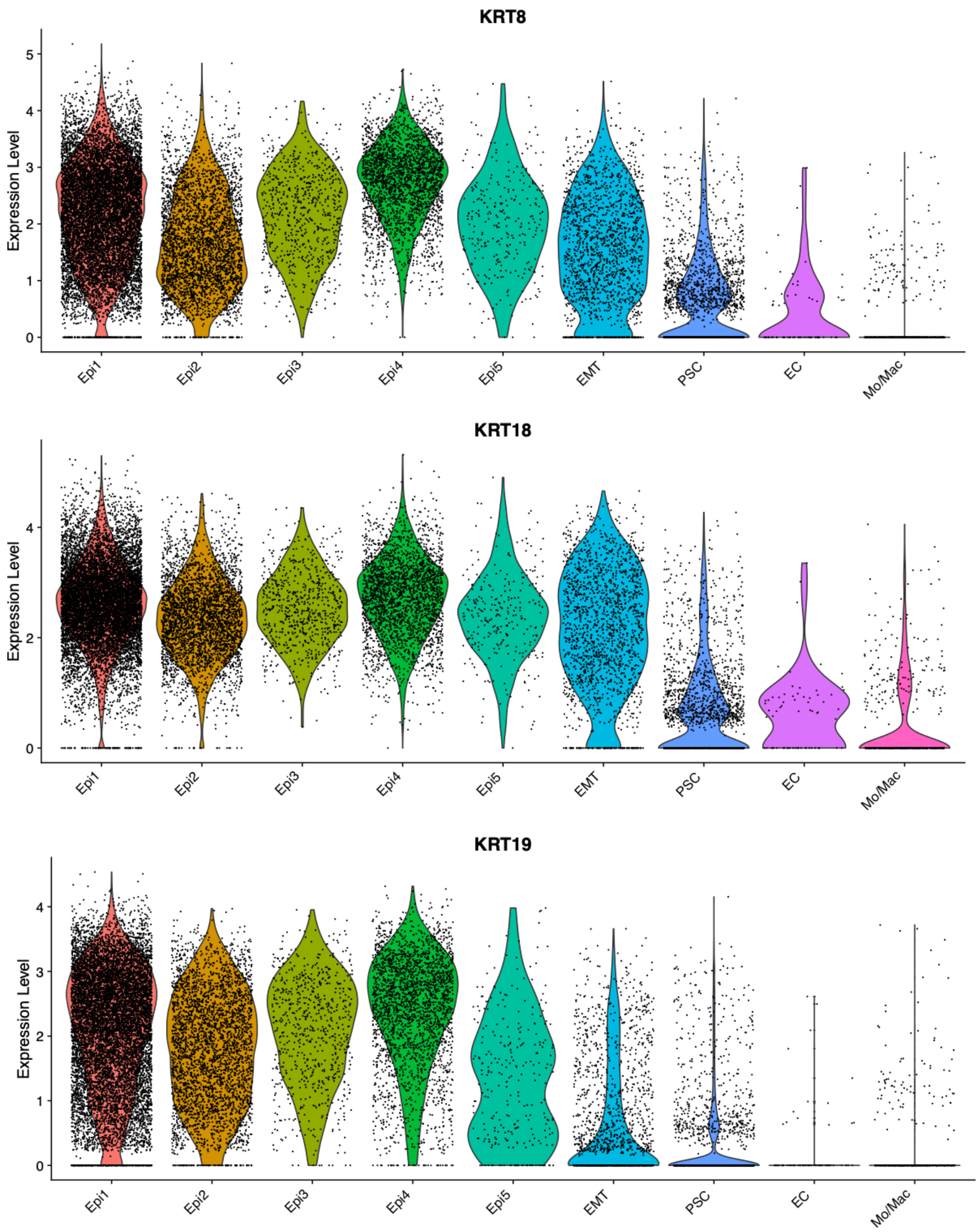


**Figure S1. Expression of keratin 8, 18 and 19 in the identified cell clusters.** Violin plot displaying keratin 8/18/19 expression of cell clusters obtained by unsupervised analysis.

### Epithelial cell composition

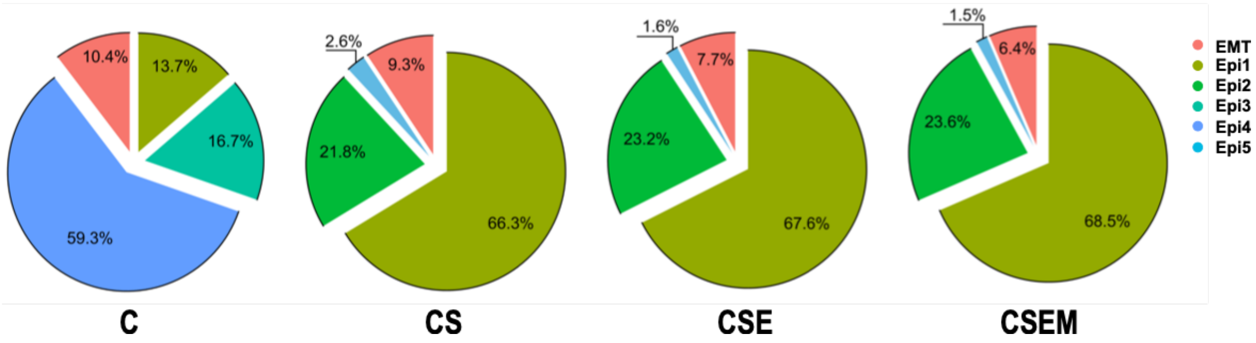


**Figure S2. Epithelial cell composition.** Pie charts showing the distribution of epithelial cell clusters as percentage of the total number of epithelial cells in spheroids with increasing levels of heterogeneity.

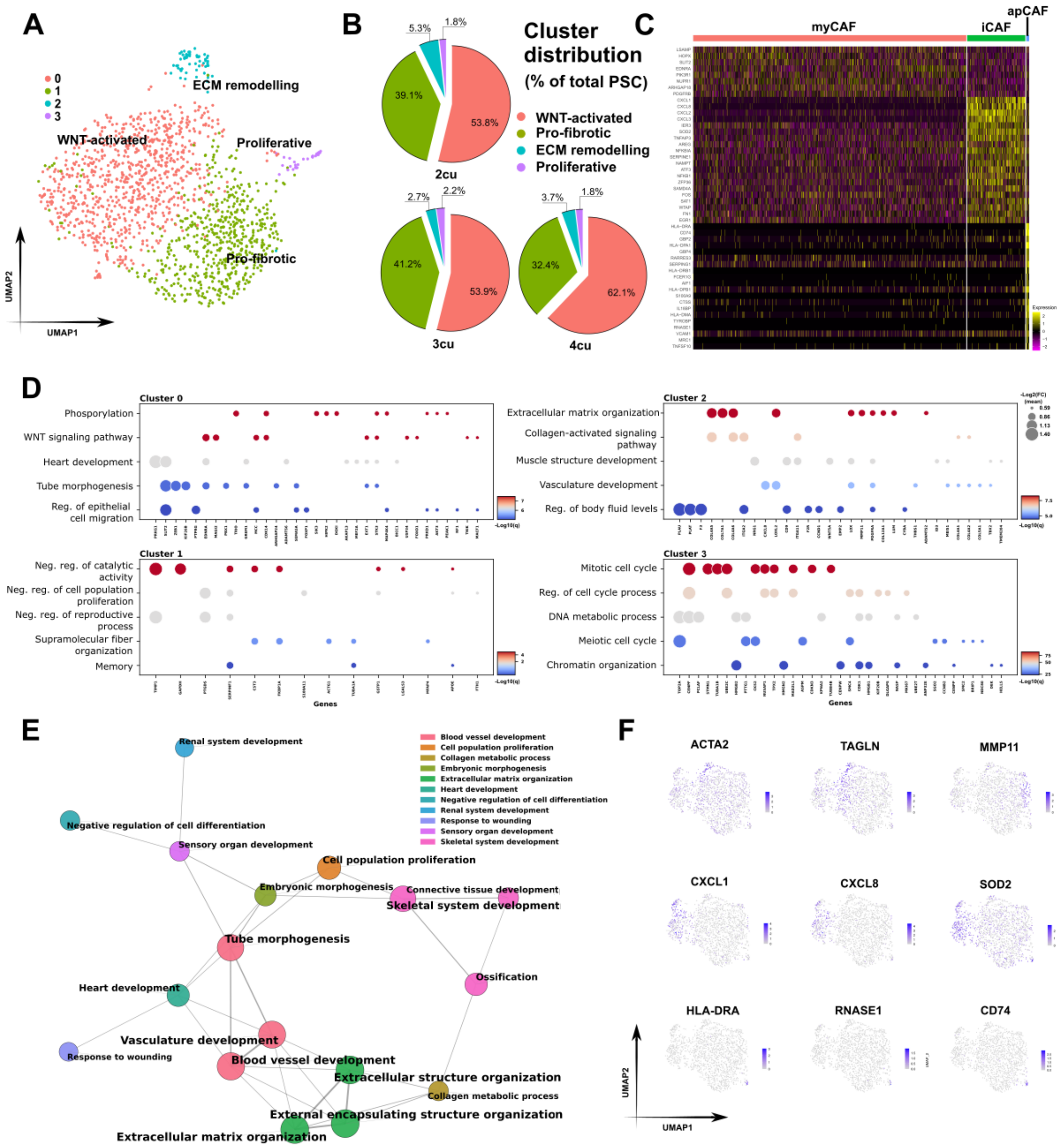


**Figure S3. PSC cells characterization.** (**A**) UMAP plot illustrating the unsupervised sub-clustering of PSC based on their gene expression profiles. (**B**) Pie charts showing the distribution of PSC clusters obtained by unsupervised analysis and displayed as percentage of the total number of PSCs in spheroids with increasing levels of heterogeneity. (**C**) Heatmap displaying the expression levels of DEGs across PSC sub-clusters following dimension reduction within the PSC cluster. (**D**) Bubble plots showing the most enriched GO Biological Process terms associated with the upregulated genes (Log2(FC) > 0.5, adj. p-value > 0.05) for each PSC cluster in heterocellular spheroids, alongside the expression of genes linked to each annotation. Color encodes −Log10(q-value) of the term and bubble size the gene expression (−Log2(FC)). (**E**) Network of the top representative Gene Ontology (GO) Biological Process (BP) terms derived from the Pathway Enrichment Analysis (PEA) performed using upregulated genes (first 200 genes, Log2(FC) > 0.5, adj. p-value > 0.05) obtained comparing PSC with the other cell clusters in all the heterogenous spheroids. Each GO BP term is represented in the network by a circle node with its size proportional to the number of input genes falling under that term, while its color represents its subset identity. The thickness of the edges connecting the terms represent the similarity score. One term from each cluster is selected to have its term description shown as label of the subset. (**F**) UMAPs showing the expression of key CAF genes. Color scale represents the expression for the gene of interest.

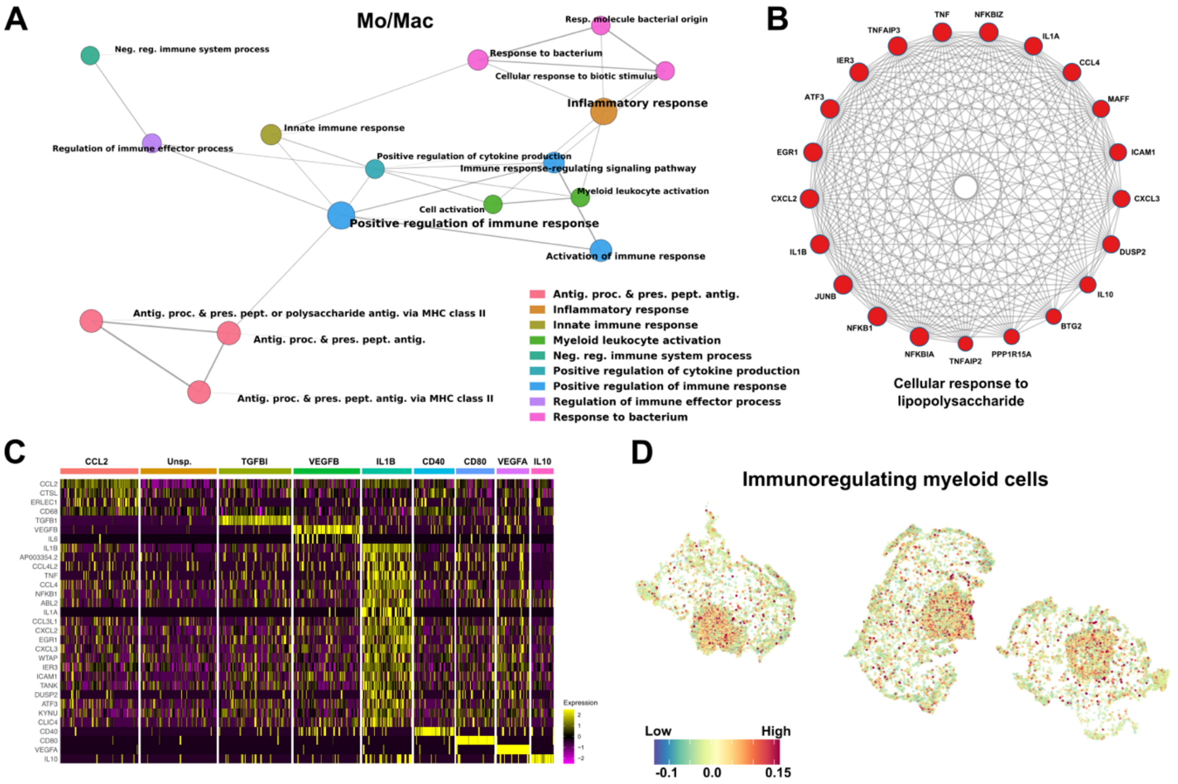


**Figure S4. Mo/Mac characterization.** (**A**) Network of the representative Gene Ontology (GO) Biological Process (BP) terms from the Pathway Enrichment Analysis (PEA) performed using upregulated genes (Log2(FC) > 0.5, adj. p-value > 0.05) obtained comparing expression of Mo/Mac cells with the other cell clusters in the spheroids. Each term is represented by a circle node. Node size is proportional to the number of input genes falling under that term. Color indicates the corresponding subset. The thickness of the edges connecting the terms represents the similarity score. One term from each cluster is selected to label the subset. (**B**) Network of highly connected upregulated genes of IL1B cluster associated with the annotation "cellular response to lipopolysaccharide" obtained with Metascape and MCODE algorithm. (**C**) Heatmap showing the expression levels of DEGs across macrophage sub-clusters in CSEM spheroids. (**D**) Spatial expression patterns of a custom gene set reporting key genes of immunoregulating myeloid cells. Colors denote program score per cell.

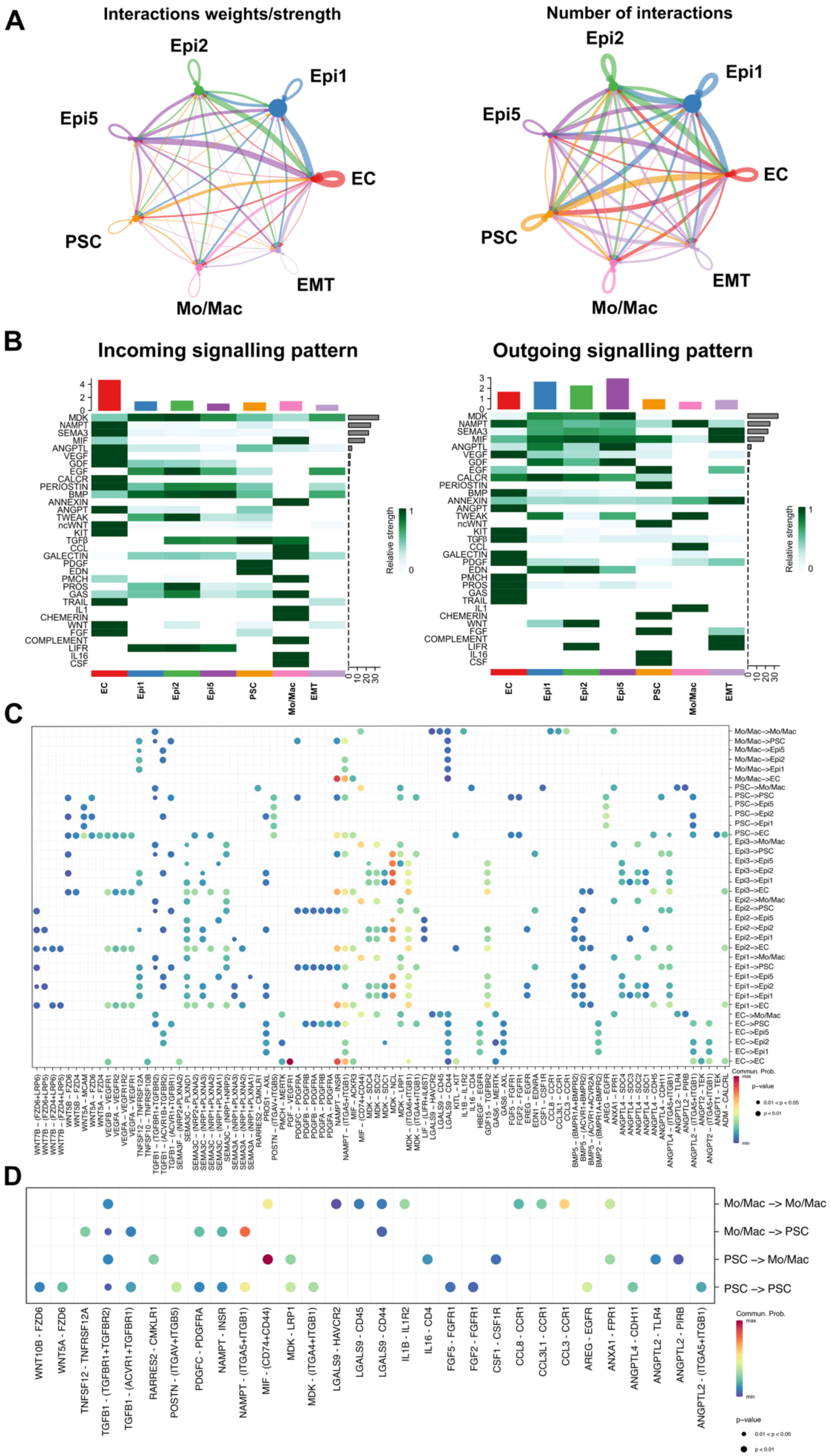
A
Interactions weights/strength
Number of interactions
Epi2
Epi1
Epi5
EC
PSC
EMT
Mo/Mac
B
Incoming signalling pattern
Outgoing signalling pattern
MDK
NAMPT
SEMA3
MIF
ANGPTL
VEGF
GDF
EGF
CALCR
PERIOSTIN
BMP
ANNEXIN
ANGPT
TWEAK
ncWNT
KIT
TGFβ
CCL
GALECTIN
PDGF
EDN
PMCH
PROS
GAS
TRAIL
IL1
CHEMERIN
WNT
FGF
COMPLEMENT
LIFR
IL16
CSF
Relative strength
EC
Epi1
Epi2
Epi5
PSC
Mo/Mac
EMT
C
Commun. Prob.
p-value
D
Mo/Mac -> Mo/Mac
Mo/Mac -> PSC
PSC -> Mo/Mac
PSC -> PSC
WNT10B - FZD6
WNT5A - FZD6
TNFSF12 - TNFRSF12A
TGFB1 - (TGFBR1+TGFBR2)
TGFB1 - (ACVR1+TGFBR1)
RARRES2 - CMKLR1
POSTN - (ITGAV+ITGB5)
PDGFC - PDGFRA
NAMPT - INSR
NAMPT - (ITGA5+ITGB1)
MIF - (CD74+CD44)
MDK - LRP1
MDK - (ITGA4+ITGB1)
LGALS9 - HAVCR2
LGALS9 - CD45
LGALS9 - CD44
IL1B - IL1R2
IL16 - CD4
FGF5 - FGFR1
FGF2 - FGFR1
CSF1 - CSF1R
CCL8 - CCR1
CCL3L1 - CCR1
CCL3 - CCR1
AREG - EGFR
ANXA1 - FPR1
ANGPTL4 - CDH11
ANGPTL2 - TLR4
ANGPTL2 - PIRB
ANGPTL2 - (ITGA5+ITGB1)
Commun. Prob.
p-value

**Figure S5. CellChat intercellular gene-based ligand-receptor interactions.** (**A**) Ribbon graphs illustrating the crosstalk between different cell types within the CSEM spheroids based on ligand-receptor interactions. Each node color represents a cell type, and the connection thickness (ribbons) indicate the strength and number of interactions between nodes. (**B**) Heatmap showing the outgoing and incoming signaling patterns between cell types in the spheroids. The heatmap indicates the relative strength of these interactions, with darker shades representing stronger interactions. The color bars at the top of the heatmap represent the sum of the signaling strength of the highlighted interactions for each cell type (column). The grey bars on the right side of the heatmap represent the sum of the strength of the interaction (row) across the different cell types.
(**C**) Dot plot illustrating the communication probabilities between various cell types in the 4cu PDAC spheroid model, focusing on ligand-receptor interactions. The size of the dots represents p-value and the color scale the probability of interactions, with warmer colors indicating higher probabilities. (**D**) Focused dot plot illustrating only the self-interactions or between PSC and Mo/Mac clusters. The size of the dots represents p-value and the color scale the probability of interactions, with warmer colors indicating higher probabilities.

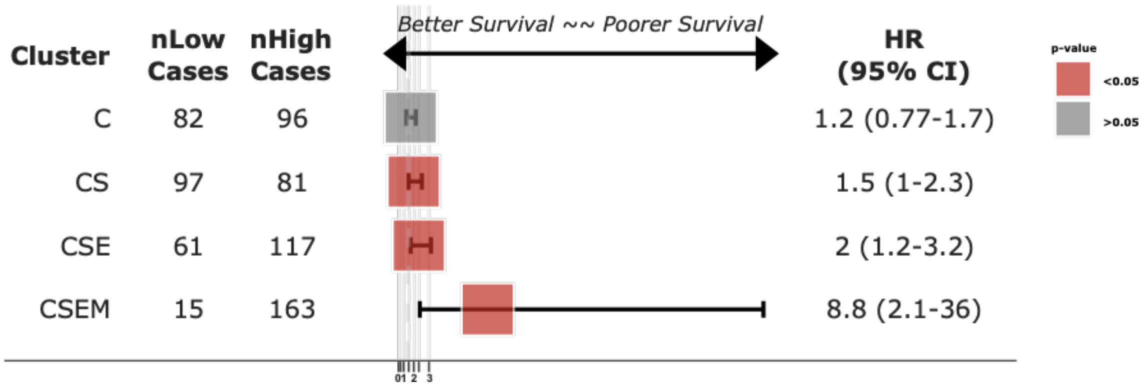


**Figure S6. Comparison of upregulated gene-associated hazard ratios for the spheroids at different heterogeneity.** Forest plot showing the hazard ratios of the up-regulated genes (Log2(FC) > 0.25, adj. p-value < 0.05) obtained by the comparison between the cells belonging to the spheroids at different composition.

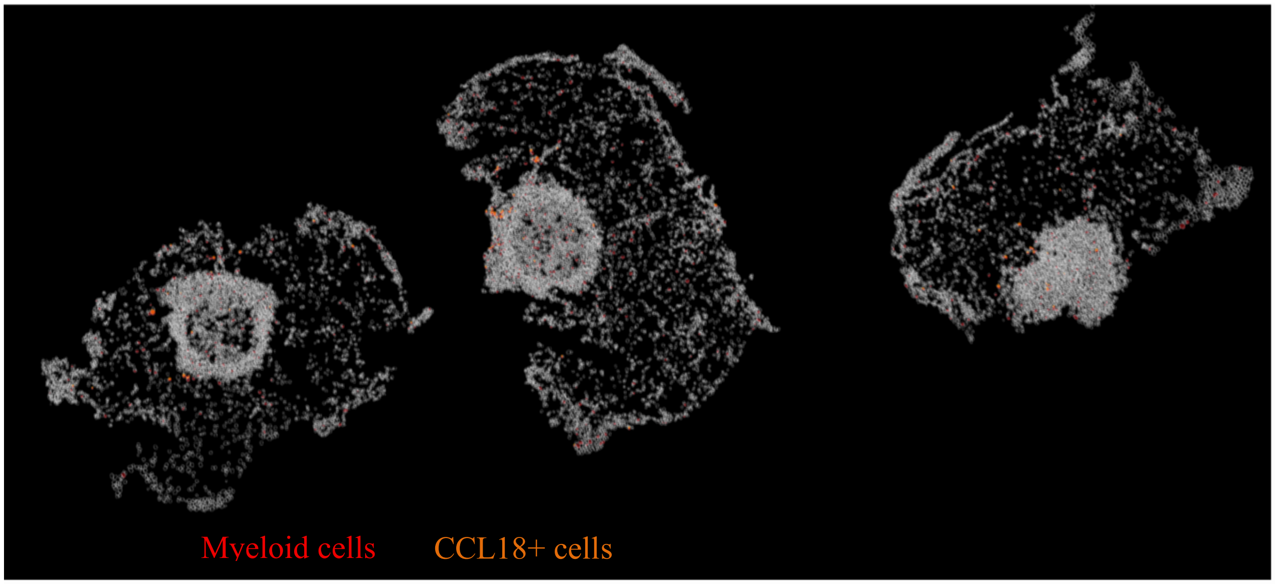


**Figure S7. Spatial distribution of CCL18 positive myeloid cells.** Myeloid cells were visualized using the expression of CD14, PTPRC, FOLR2, HLA-DRA, HLA-DRB1, HLA-DRB9, HLA-DRB6, HLA-DRB6, HLA-DRB5, MRC1, CD163, MARCO, VSIG4, C1QA/B/C genes.

**Table S1. CAF markers for dimensional reduction.** Markers used for dimensional reduction of scRNA-seq data for PSC cluster. The table lists phenotype-specific genes associated with myCAFs, iCAFs and apCAFs, which were used to distinguish these subtypes.

| Phenotype | Markers |
|---|---|
| **myCAF** | *CST1, APOD, MGP, MMP11, TAGLN, POSTN, ACTA2, COL1A1, COL1A2, COL11A1, COL7A1, CTHRC1, SFRP2, SPARC, ASPN* |
| **iCAF** | *MMP3, CXCL14, IL24, CXCL8, CXCL5, MMP1, ADM, CXCL2, IER3, IL11, CXCL16, CXCL13, FTH1, CXCL3, SOD2* |
| **apCAF** | *IGFBP5, CD74, CCN3, HLA-DRA, HLA-DRB1, HLA-DRB5, HLA-DQA1, HLA-DPB1, TCF4, HSPD1, RNASE1, PLD3, S100B, ITGB8, LAPTM5* |

**Table S2. Macrophage markers for dimensional reduction.** Markers used for dimensional reduction of scRNA-seq data for Mo/Mac cluster. The table lists phenotype-specific genes associated with M1 and M2 macrophages, which were used to distinguish these subtypes.

| Phenotype | Markers |
|---|---|
| **M1** | *CD14, CD68, CD40, CD80, CD86, TNF, IL6, IFNG, IL1B, NOS* |
| **M2** | *CD14, CD68, CD206 (MRC1), CD163, TGFBI, IL4, IL10, VEGF, CCL2, MMP* |

### Extended methods

#### LC–MS/MS analyses

Peptides were first resolved at a constant flow rate of 300 nl/min using 0 to 30% solvent over 148 min, raised to 48% over 16 min, then ramped to 100% in 8 min and held for 8 min with a flow rate of 400 nl/min. Eluting peptides were online injected into the mass spectrometer with electrospray ionization performed at a 2.1 kV static spray voltage; the temperature of the ion transfer tube was set to 300 °C, and the RF lens voltage was set to 55%. Full scan MS spectra were acquired at a resolution of 60,000 within the m/z range of 350-1550, accumulating to 'Standard' pre-set automated gain control (AGC) target. Multiply charged ions were selected for further fragmentation with higher energy collision dissociation (HCD) performed at 32% normalized collision energy (NCE). Scans were performed at a resolution of 7500 with a dynamic exclusion time of 30 sec and a 1 m/z isolation window.

#### Database Search and Analysis

Raw files were searched against the human UniProt database (April 2024 with 204093 entries) including common contaminants using MaxQuant software (version 2.5.2.0) with the integrated Andromeda search engine. Digestion was set as Trypsin/P with a maximum of 2 missed cleavages allowed. Fixed modification was set as cysteine carbamidomethylation, and variable modifications as protein N-terminal acetylation and methionine oxidation. The match-between-run and label free quantification (LFQ) features were enabled for identification and quantification. A false discovery rate (FDR) of 1% was applied to both peptide spectrum matches (PSMs) and protein identification using a target-decoy approach.

Data filtering was performed using Perseus software (version 2.0.11). Proteins only identified by site, potential contaminants and reverse peptides were removed. LFQ intensities were $Log_2$ transformed and proteins that were quantifiable in at least three out of four replicates were retained after imputation based on normal distribution. To filter for significant changes between the samples, two-sample unpaired Student's t-test was performed. FDR-corrected p-values (q-values) were calculated from 250 randomizations and considered significant if they were lower than 0.05. The data was visualized with GraphPad Prism 10.

#### Stereo-seq raw data processing (Alignment and UMI counting)

Two core tools in SAW accomplish these steps. We utilized the SAW software's mapping tool to align read1's CIDs with the Stereo-seq chip's coordinates, allowing a single base mismatch. This resulted in "Valid CID Reads," which were then supplemented with coordinate data. The subsequent "Clean Reads" were obtained by filtering out MIDs with polyA sequences and low-quality bases (N or more than two bases with a score below 10), as well as mRNA with polyA. The Clean reads were aligned to the human reference genome GRCh38 release 93 (GRCh38-3.0.0 reference package downloaded from 10x Genomics), using STAR (v2.5.3) [1] and the number of reads aligned to regions such as exons, introns, and intergenic regions were counted according to the gene annotation files. Using Bam2Gem (https://github.com/BGIResearch/handleBam), the corresponding relationships between the unique mapping reads aligned to the reference genome and the genes were determined, and the expression levels of all genes were calculated according to MID correction. Through quantification of gene expression, the final output was the expression matrix of all genes detected in the tissue section, which was stored in a GEM format file. The bin1 represents one spot with 220 nm diameter in section, and bin n represents combined bins in an N × N square area.

**Metascape**

For pathway enrichment or gene set enrichment analysis (respectively PEA and GSEA) differentially expressed genes (upregulated, downregulated or both) were processed using Metascape [106]. Custom PEA and GSEA was performed using the following settings: minimum overlap: 3, p-value cutoff 0.05, minimum enrichment 1.5. Using these settings Gene Ontology Biological Processes (GO BP), KEGG pathway and Hallmark gene set databases were used to determine the relevant annotations. Protein-protein interaction (PPI) analysis was performed using Metascape using either physical core or combined core. All protein-protein interactions among input genes were extracted from PPI data source and formed a PPI network. GO enrichment analysis was applied to the network to extract biological meanings. MCODE algorithm was then applied to this network to identify neighborhoods where proteins are densely connected. Part of the image generation was performed using Phyton 3. Phyton scripts were drafted with the assistance of AI tools (ChatGPT, Open AI).

**Image processing and cell segmentation**

First, the total number of genes under a single DNB was transformed into an image and registered with the single-stranded DNA staining image using ImageJ [2] software. The Python package StereoCell [3] was used to perform nuclear segmentation. The gaussian mixture model (GMM) implemented in StereoCell was used to fit the molecular distribution under a cell-nucleus mask. The probability of extracellular molecules with a fitting range (100 × 100 pixels) was then calculated according to the GMM, and the pixels outside the mask were resegmented to obtain the complete cell boundary. Unique molecular identifiers from all DNBs in the same putative single cell (referred to as a cellbin) were aggregated to generate a gene expression matrix for downstream analysis.

**Supplementary video 1 (separate file)**

Representative video showing neutrophils (in blue) injected in the vasculature (in orange) within the OrganiX$^{TM}$ and interacting with cancer cells (in green).

**Data file S1. (separate excel file)**

Differential gene expression of the identified spheroid cell clusters

**Data file S2. (separate excel file)**

Pathway enrichment analysis for upregulated genes of epithelial clusters

**Data file S3. (separate excel file)**

Differential gene expression of EMT cells of heterocellular spheroids compared to C spheroids

**Data file S4. (separate excel file)**

Differential gene expression of PSC cells of CSE and CSEM spheroids compared to CS spheroids
